# Sequence-dependent binding modes of INO80 control +1 nucleosome positioning

**DOI:** 10.64898/2026.08.12.744421

**Authors:** Mariia Likhodeeva, Annika Brem, Alberto López-Francos López-Romero, Drin Shabani, Franziska Därr, Dirk Kostrewa, Katja Lammens, Manuela Moldt, Olga Fettscher, Blaine Bartholomew, Philipp Korber, Karl-Peter Hopfner

**Author notes:** Correspondence: Prof. Dr. Karl-Peter Hopfner, Gene Center, Ludwig-Maximilians-Universität Munich Feodor-Lynen-Str. 25, 81377 Munich, Germany. These authors contributed equally: Mariia Likhodeeva, Annika Brem.

## Abstract

Cellular self-organisation counteracts entropy at the expenditure of energy. In case of the first level of nuclear DNA organisation, this relates to regular nucleosome arrays and interspaced nucleosome-depleted regions (NDRs), for example at promoters or replication origins^1–5^. The organisation of nucleosomes as building blocks of chromatin is orchestrated by the collective activities of ATP-dependent chromatin remodellers^6–9^. Yet, how remodellers achieve positional specificity, in particular regarding the promoter-proximal +1 nucleosomes, remains unclear. Here, we show that the *S. cerevisiae* chromatin remodeller INO80 unexpectedly distinguishes DNA sequence asymmetry within the +1 nucleosome of the *SWH1* gene through distinct inhibited and active nucleosome-binding modes. In structural and biochemical analyses of INO80 on nucleosomes with the endogenous sequence, we identified an inhibited binding mode where the entire INO80 remodelling unit flipped on the +1 nucleosome. INO80 adopted this remodelling-incompetent binding mode when facing the promoter, but a remodelling-competent mode when facing the gene body. This directional read-out of intra-nucleosomal DNA sequence asymmetry, together with extranucleosomal NDR sequence features, prevented nucleosome sliding into the NDR while permitting array formation over the gene. Our work shows how DNA features contribute to ATP-dependent self-organisation of promoter chromatin by INO80.

---

Eukaryotic nuclear DNA exists in a highly organised nucleoprotein complex termed chromatin. The basic building block of chromatin is the nucleosome, ∼147 base pairs (bp) of DNA wrapped around histone protein octamers. A principle of chromatin organisation is the presence of nucleosome-depleted regions (NDRs) flanked by arrays of regularly spaced nucleosomes. The arrays are typically found across gene bodies, whereas NDRs demarcate regulatory regions such as promoters, enhancers and replication origins^1–5^.

The nucleosomes flanking NDRs are designated +1 in the downstream and −1 in the upstream direction of transcription, and subsequent array nucleosomes are denoted +2, +3, … or −2, −3, …, respectively. In *Saccharomyces cerevisiae* (*S. cerevisiae* or *Sc*), +1 nucleosomes often occupy well-defined positions. They co-define the transcription start site (TSS), play key roles in transcription initiation and origin firing^10,11^ and “phase” adjacent nucleosomal arrays^4,12–14^. Such phased arrays repress cryptic transcription^15–18^ and assist origin firing^19^.

The organisation of nucleosomes around NDRs and in genic arrays is a prime example of energy-driven cellular self-organisation and orchestrated, among others, by the collective activities of ATP-dependent chromatin remodellers^6–8,20,21^. Remodellers couple ATP hydrolysis to nucleosome sliding, histone eviction and histone variant exchange. Remodeller-intrinsic mechanisms together with regulatory cues shape the nucleosomal landscape^7^. Notably, in some cases, just the DNA sequence alone may direct a remodeller to generate defined nucleosome positions.

The *Sc*INO80 complex^22^ is the leading example for this sequence feature-linked mechanism as it can position +1 and -1 nucleosomes on its own across the yeast genome *in vitro*^6,23^. INO80 is organised into a modular structure with a nucleosome-sliding C- and regulatory N- and A-modules^24–28^ (Fig. 1A). Like its structural relative SWR1, INO80 is activated by the presence of long extranucleosomal DNA (>30 bp^29,30^), which is bound by INO80’s A- and N-modules^31–35^. However, the activation of INO80 by extranucleosomal DNA generates a mechanistic paradox regarding INO80’s +1 nucleosome positioning activity. NDRs are sufficiently long to accommodate INO80’s regulatory modules, and their often dA/dT-rich nature should in particular allow A-module binding^3,36^. Thus, INO80 should slide nucleosomes into the NDR, rather than stop and position them at the boundary of the NDR. A possible key to resolving this paradox may be that not only DNA length, but also DNA sequence features, both intra- and extranucleosomal, affect nucleosome positioning^6,23,37^, like intranucleosomal DNA rigidity in case of remodelling by INO80^38^ or Chd1^39^. In this study, we use reconstituted mononucleosomes and nucleosomal arrays around the +1 nucleosome of the *S. cerevisiae SWH1* gene to uncover the interaction of INO80 with nucleosomal arrays assembled on native promoter-flanking DNA sequences and reveal the mechanism of DNA sequence-guided +1 nucleosome positioning by INO80.

**Fig. 1:**
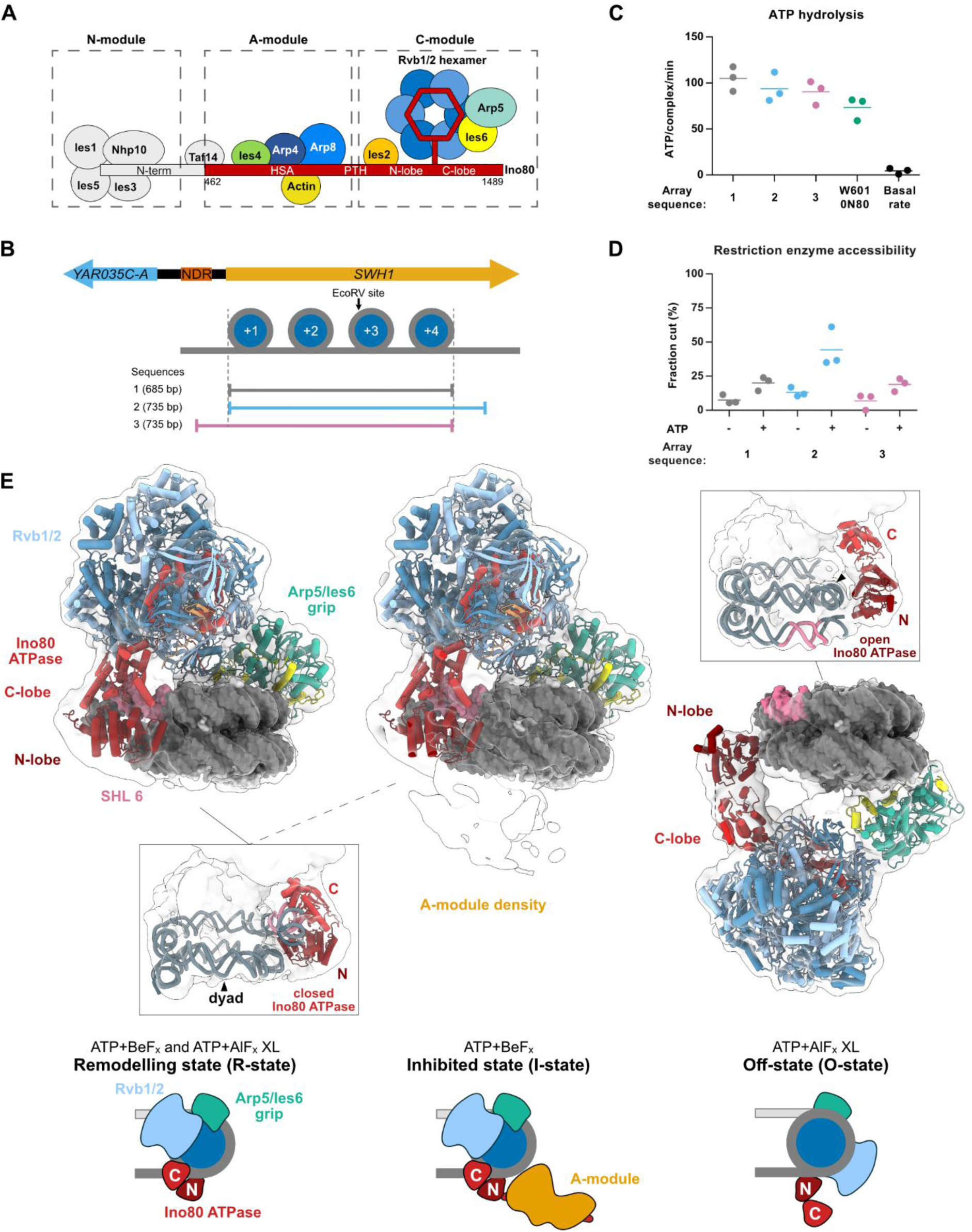
INO80 adopts different nucleosome-binding modes on promoter-proximal nucleosomal arrays. **A,** *Sc*INO80 modular organisation. *Sc*Ino80^ΔN^ truncation leaving only amino acids 462-1489 are indicated. HSA, Helicase SANT-associated; PTH, post-HSA. **B,** Choice of the natural array sequences from the *SWH1* gene with indicated EcoRV site. NDR, nucleosome-depleted region. **C,** ATP hydrolysis rates of INO80^ΔN^ on the arrays and a control Widom 601 mononucleosome with 80 bp overhang of synthetic DNA (n = 3). **D,** Quantification of the fraction of EcoRV-digested array DNA. In the restriction enzyme accessibility (REA) assay, arrays were incubated with EcoRV and INO80^ΔN^ in absence or presence of ATP (n = 3). **E,** Cryo-EM density maps (low-pass filtered to 10 Å) with fitted models of different INO80 binding modes (R-state, I-state and O-state) on the nucleosomal arrays with the representative cartoons. The difference in open and closed Ino80 ATPase binding is visualised in the frames. SHL 6, ATPase binding site in the R-state, is marked in pink. SHL, superhelical location; XL, crosslinked.

## INO80 adopts different nucleosome-binding modes on promoter-proximal nucleosomal arrays

We chose endogenous sequences based on the *S. cerevisiae SWH1* gene, since synthetic Widom 601 sequences, used in most structural studies, do not lead to proper array formation *in vivo*^40,41^. The *SWH1* +1 nucleosome sequence combines strong intrinsic positioning capability by salt gradient dialysis (SGD) *in vitro* with being a *bona fide* well-positioned +1 nucleosome *in vivo*^42^. We tested three sequences (1-3) encompassing the +1 to +4 nucleosomes of *SWH1* (Fig. 1B). Tetra-nucleosomal arrays assembled well on these sequences in SGD (Fig. S1A). The nucleosomal arrays entailed approximately 30 bp linker DNAs, which correspond to the minimal linker length for remodelling by INO80^30^. All arrays induced robust ATPase activity in INO80^ΔN^ (lack of the N-module’s Nhp10 subunit had no effect on +1 positions *in vivo*^43^, Fig. 1C, D). Array 1 showed modest remodelling, as expected from the short linkers and our restriction enzyme accessibility assay may not efficiently pick up local remodelling due to the natural cleavage site occurring within the +3 nucleosome. Array 2, with a linker extending into the gene body, showed moderately increased remodelling, probably caused by the stimulatory effect of additional extranucleosomal DNA on nucleosome sliding^32^. Array 3, extending into the promoter, showed similar remodelling as array 1, suggesting a lack of sliding toward the NDR (Fig. 1D, Fig. S1B). In summary, all sequences form homogeneous arrays and robustly stimulate INO80’s ATPase, with modest remodelling aimed at helping us to identify both sliding-poised and sliding-inhibited conformations.

We used array 1 for interrogating the interaction of INO80 bound to promoter-proximal chromatin structures. To reduce nucleosomal stacking, we introduced the H4K16Q mutation that mimics the “open” chromatin achieved with H4K16ac^44,45^. INO80^ΔN^ slid a mononucleosome with H4K16Q as efficiently as with wild type H4 (Fig. S1C), consistent with prior observations that H4 tails do not *per se* affect INO80 activity^30^. We recorded two cryo-EM datasets of INO80^ΔN^ bound to array 1 in the presence of ATP and BeF_x_ or AlF_x_. In the ATP+BeF_x_ dataset, we observed the previously obtained canonical nucleosome-binding mode, with Ino80’s ATPase “motor” engaged with unwrapped entry DNA at SHL 6 and the Arp5/Ies6 “grip” bound to SHL 3, poised to pump DNA into the nucleosome core particle (NCP)^24,26^. The A-module was either not visible in this mode or was folded back in recently found autoinhibited conformation^33^, consistent with the effect of short linkers. We denote these modes as R-state (“remodelling”, extended A-module) and I-state (“inhibited”, folded back A-module) (Fig. 1E, Fig. S2).

Intriguingly, in the ATP+AlF_x_ dataset, we observed along R- and I-states a new nucleosome-binding mode that we called the O-state (“off-state”). The O-state was less stable, as its presence could be increased by mild glutaraldehyde crosslinking (Fig. S3). It strongly deviated from all other conformations observed for INO80 in that the Ino80 ATPase was not placed at SHL 6, but at SHL 1, and Arp5/Ies6 was bound to SHL 5 instead of SHL 3. This binding mode is different from the SHL 2-bound conformations of other remodellers^7^, and notably, the two lobes of the Ino80 ATPase were open (Fig. 1E). This indicates a defunct ATPase, even in presence of ATP+AlF_x_.

These results established that the arrays with endogenous sequence captured a range of different binding modes of INO80: either the Ino80 ATPase was bound at SHL 6 and poised for sliding (R- and I-states), or at SHL 1 and incapable of ATP hydrolysis and sliding (new O-state).

## Promoter-facing INO80 binds the SWH1 +1 nucleosome in the off-state

In our structural analysis of arrays, we could not distinguish to which of the four nucleosomes INO80^ΔN^ was bound in which conformations. However, given INO80’s limited remodelling of arrays towards the NDR and the long-standing question of how INO80 determines +1 nucleosome positioning (Fig. 1B-D), we tested whether the O-state could be arising on the +1 nucleosome and lead to setting this position. We assembled a mononucleosome on DNA comprising the *SWH1* +1 nucleosome and 80 bp upstream promoter sequence (denoted 80S0, Fig. 2A). This sequence assembled a homogeneous nucleosome as judged by native gel electrophoresis (Fig. 3C, 80S0 -ATP lane). The cryo-EM structure of 80S0 in complex with INO80^ΔN^ in presence of ATP+AlF_x_ (or BeF_x_ with similar results) revealed that most, if not all, nucleosome-bound INO80 was in the O-state (even without crosslinking), although the 80 bp extranucleosomal promoter would allow INO80 to adopt a promoter-facing R-state. Again, the Ino80 ATPase was in an open conformation, with N- and C-lobes widely separated even in presence of ATP/ADP+AlF_x_/BeF_x_ (Fig. 2A). Only the N-lobe was bound at SHL 1, while the C-lobe remained distant through its tight interaction with the Rvb1/2 subcomplex.

**Fig. 2:**
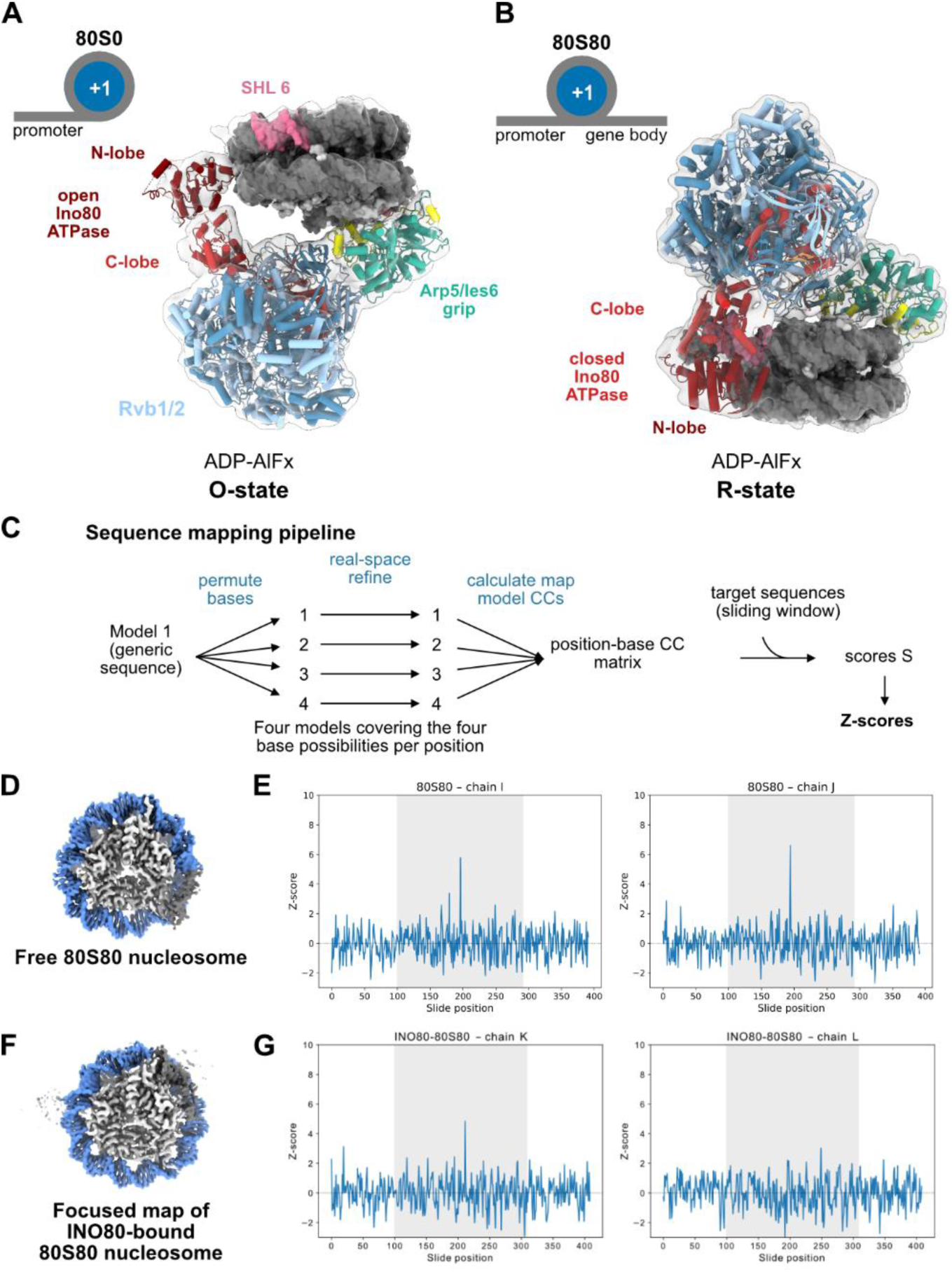
INO80 distinguishes *SWH1* +1 nucleosome asymmetry through different binding modes. **A,** Cryo-EM density map (low-pass filtered to 10 Å) with fitted model. Promoter-facing INO80 binds the *SWH1* +1 80S0 nucleosome in the O-state. SHL 6 is marked in pink. **B,** Cryo-EM density map (low-pass filtered to 10 Å) with fitted model. Gene body-facing INO80 binds the *SWH1* +1 80S80 nucleosome in the R-state. **C,** Nucleosomal sequence mapping pipeline (see Suppl. Method). **D,** Cryo-EM density map of a free 80S80 nucleosome. **E,** Z-scores of 80S80 forward strand correlated with both nucleic acid chains (I and J) in the cryo-EM map of the free 80S80 nucleosome. **F,** Cryo-EM density focused map of an INO80-bound 80S80 nucleosome. **G,** Z-scores of 80S80 forward strand correlated with both nucleic acid chains (K and L) in the cryo-EM map of the INO80 bound 80S80 nucleosome.

**Fig. 3:**
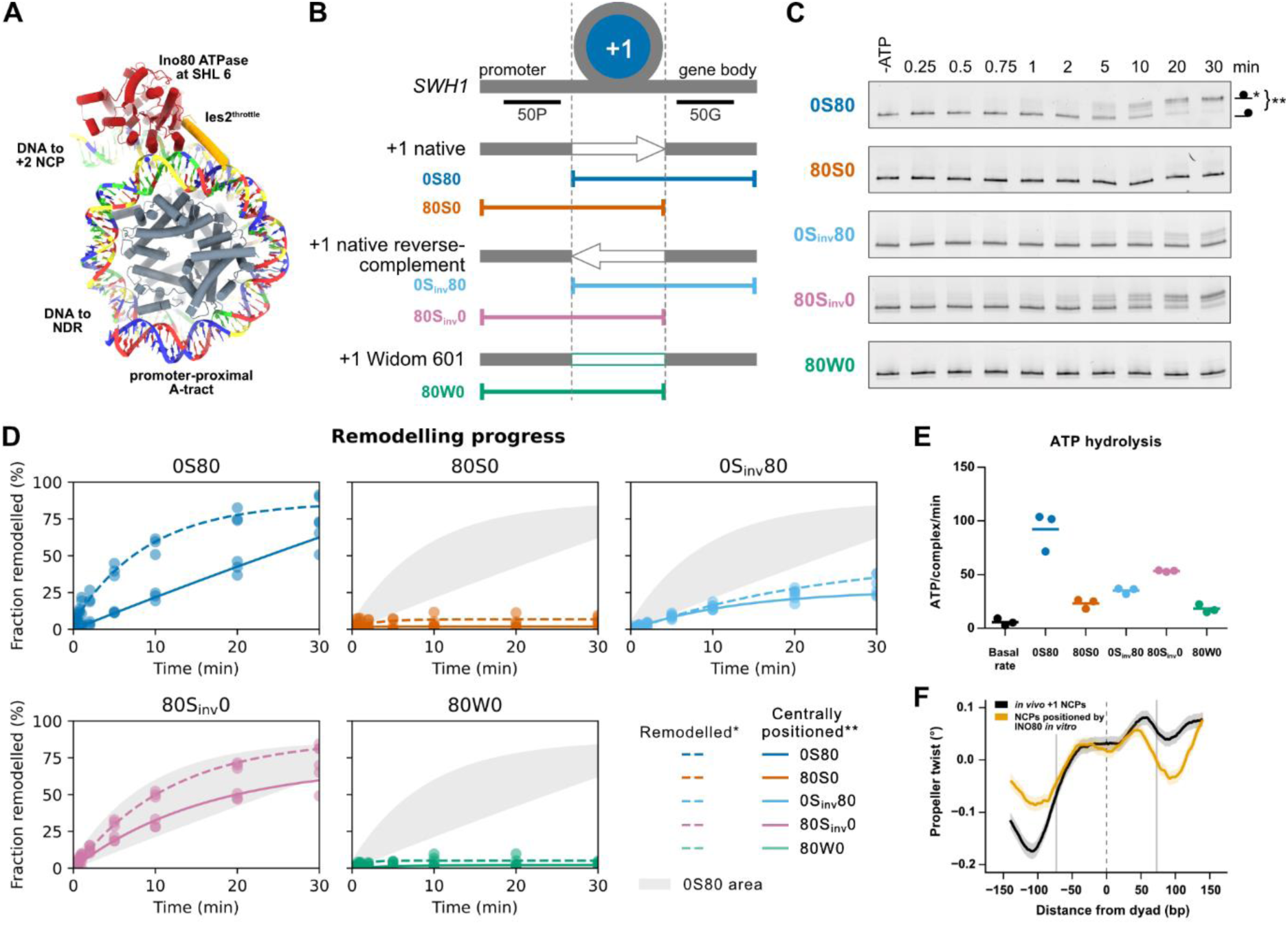
INO80 slides the highly asymmetric *SWH1* +1 nucleosome away from, but not into, the promoter. **A,** Model of an INO80-bound 80S80 nucleosome with mapped DNA sequence. dA, dT, dG, and dC nucleotides are coloured red, blue, green, and yellow, respectively. **B,** Architecture of nucleosomal substrates assembled on the following sequences: natural *SWH1*, reverse complement *SWH1* +1 nucleosome, and Widom 601 replacing *SWH1* +1 nucleosome, with extranucleosomal DNA reaching either into the promoter or gene body sequences. 50P and 50G demarcate the DNA sequences used for measuring A-module binding (see Fig. S8A). **C,** Representative sliding assay native gels for the above-mentioned nucleosomal substrates (see B). **D,** Remodelling progress of each nucleosomal substrate. In each panel, the upper dashed line fit shows the total remodelling products (all but end-positioned nucleosomes, marked with *), the lower solid line fit shows the emergence of the centrally positioned nucleosomes (marked with **). **E,** ATP hydrolysis rates of the nucleosomal substrates as in panel B. **F,** Composite profiles for all *in vivo* +1 nucleosomes (black trace) and for all nucleosomes positioned by INO80 *in vitro* (orange trace) show a similar propeller twist gradient. Vertical dashed line marks the dyad, vertical solid lines mark the nucleosome borders (+/- 73 bp from the dyad).

In summary, the *SWH1* +1 nucleosome sequence strongly suppressed an “into the NDR”- sliding R-state in favour of a new O-state with a defunct Ino80 ATPase.

## INO80 distinguishes SWH1 +1 nucleosome asymmetry through different binding modes

To test whether the *SWH1* +1 nucleosome prevents formation of active INO80 states *per se*, or only in the direction of the promoter, we tested an 80S80 nucleosome, encompassing the *SWH1* +1 nucleosome sequence and 80 bp flanking DNA from both the promoter and gene body sides. Here, the INO80 A-module could bind either of the extranucleosomal DNAs. Structure determination revealed that 80S80-bound full-length INO80 (INO80^fl^) was now in the canonical active R-state with N- and C-lobes properly closed around SHL 6 and Arp5/Ies6 bound to SHL 3 (Fig. 2B).

These data indicated that the *SWH1* +1 nucleosome does not intrinsically prevent formation of an active sliding state of INO80 but may do so specifically in the promoter-facing orientation. To test this, we sought to determine whether the R-state on 80S80 is specifically arising in the gene-body facing orientation. We developed an approach to score DNA sequences in cryo-EM maps based on map-model correlations in a sliding-window approach (Suppl. Method, Fig. 2C). Analysis of a free 80S80 nucleosome (Fig. 2D) revealed two high Z-score matches corresponding to the same sequence in opposite orientations (Fig. 2E). This is due to a cryo-EM processing drawback: the two pseudo-symmetric nucleosome orientations often cannot be distinguished, which results in an averaged map with both orientations superimposed. Importantly, our approach identified the previously mapped *SWH1* +1 position^42^ with single-base resolution and could even deconvolute two superimposed DNA sequences.

To reveal whether INO80 bound to 80S80 facing the promoter, the gene-body, or either, we derived a 2.75 Å map of the INO80-bound 80S80 nucleosome using focused refinement (Fig. 2F). In this case, we got only a single high Z-score corresponding to the gene-body facing R-state (Fig. 2G). Altogether, these data revealed that the binding mode of INO80 on the *SWH1* +1 nucleosome was strikingly dependent on the nucleosome orientation, with promoter-facing orientations promoting an O-state and gene body-facing orientations promoting an R-state. Furthermore, of the two possible binding conformations, the gene body-facing R-state predominated, which concurs with the O-state being less stable.

## INO80 slides the highly asymmetric SWH1 +1 nucleosome away from, but not into, the promoter

To validate the structural observations, we performed sliding assays with the *SWH1* +1 nucleosome, with sliding directionality enforced via the presence of extranucleosomal DNA only on the promoter or the gene body side (80S0 versus 0S80, Fig. 3B). INO80^ΔN^ could robustly slide 0S80 into the gene body sequence (Fig. 3C, D) but not 80S0 into the promoter sequence (Fig. 3C, D). Furthermore, the ATPase rate was high on 0S80, similar as on a Widom 601 0N80 mononucleosome (Fig. 1C, W601 0N80), but low on 80S0 (Fig. 3E). Altogether, these data are consistent with the structural findings that INO80 binds 0S80 in a sliding-competent R-state, while 80S0 enforces a sliding- and ATP hydrolysis-inhibited O-state.

To dissect the roles of nucleosome orientation versus flanking extranucleosomal DNA, we analysed INO80^ΔN^ sliding and ATP hydrolysis on nucleosome variants comprising modified intranucleosomal DNA segments (Fig. 3B). We inverted the *SWH1* +1 nucleosome sequence with respect to extranucleosomal promoter and gene body DNAs (S_inv_) and also swapped it with a Widom 601 sequence (W). Intriguingly, 80S_inv_0 showed reasonable remodelling into the promoter region, in contrast to 80S0 (Fig. 3C, D), indicating that intranucleosomal DNA features of *SWH1* +1 nucleosome strongly dictate the INO80 remodelling directionality. It is of note that 80S_inv_0 displayed a higher amount of intermediate nucleosome species (Fig. 3C). Such longer-lived intermediates could be the result of the natural A-tracts in *SWH1* promoter DNA consistent with sliding experiments on engineered extranucleosomal A-tracts^36^ (Fig. 3D). 0S_inv_80 displayed substantially reduced but detectable remodelling, while 80W0 even showed arrested sliding and low ATP hydrolysis rate. Thus, gene body extranucleosomal DNA can, to a low amount, rescue the repressed upstream sliding of *SWH1* +1, while *SWH1* promoter DNA strongly inhibits sliding of a Widom 601 nucleosome (Fig. 3C, D). Thus, overall sliding capability results from both extra- and intranucleosomal features, with a possible context-dependent domination of one or the other.

Our data show that the *SWH1* +1 nucleosome is highly asymmetric with respect to INO80 binding and sliding competence. A potential intranucleosomal feature promoting this striking asymmetry is an A-tract at the NDR-facing SHL 6/7 (Fig. 3A). A-tracts feature prominently in DNA shape prediction plots, e.g., as low propeller twist, and their increased rigidity can impair remodelling speed^42^. In the case of *SWH1* +1 nucleosome, the A-tract at SHL 6/7 might prevent the Ino80 ATPase from gripping DNA to adopt a stable R-state. Consistently, we found an enrichment of long A-tracts on the NDR-facing side compared with the gene body-facing side in +1 nucleosomes in *S. cerevisiae* (Fig. S4). More generally, a skewed asymmetry with respect to propeller twist is not only found for the sequence of the *SWH1* +1 nucleosome but typical for all *in vivo* +1 nucleosome sequences and for the sequences in all nucleosomes that are positioned by INO80 on its own in a genome-wide nucleosome positioning assay (Figs. 3F, S5-7).

## INO80 integrates nucleosomal and NDR sequence features into sliding arrest

The strong intranucleosomal contribution to the asymmetry of *SWH1* +1 nucleosome sliding by INO80 raised the question of which role the A-module plays in the promoter-facing O-state. Binding of >30-40 bp linker DNA by the A-module is a prerequisite for robust INO80 activation^30,32,33,46^. Thus, we first tested whether A-module can bind the *SWH1* NDR DNA and whether this potential extranucleosomal DNA binding can occur in the O-state. Promoter DNA showed a 3-fold reduced binding compared to gene body DNA (Fig. S8A, Table S1). The reduced but still substantial affinity may contribute to the moderate inhibitory effect in promoting sliding of *SWH1* +1 nucleosome to the central position (0S80 vs 80S_inv_0 in Fig. 3C, D), but may not stop sliding into the NDR simply by dissociation of the A-module. Likewise, robust sliding of 80S_inv_0 suggests that the A-module can bind the NDR at least in the R-state when sliding a nucleosome from the gene body towards the promoter.

In prior work, we investigated various Widom 601-derived sequences to study the effect of intranucleosomal DNA rigidity on sliding by INO80^38^. One of the studied sequences (denoted M2_rev_) assembled into multiple nucleosomal species with different extranucleosomal DNA lengths. Cryo-EM analysis of this population in complex with INO80^fl^ uncovered an intriguing compatibility between upstream extranucleosomal DNA A-module binding and the O-state. In the M2_rev_ dataset, we find the R-state (not shown) as major species along with the O-state as a minor species (Fig. 4A). Notably, the nucleosome of the O-state showed ∼20–25 bp of extranucleosomal DNA on one side and ∼50–55 bp of entry DNA on the other side potentially stabilising the overall complex including the A-module (Fig. S8B).

**Fig. 4:**
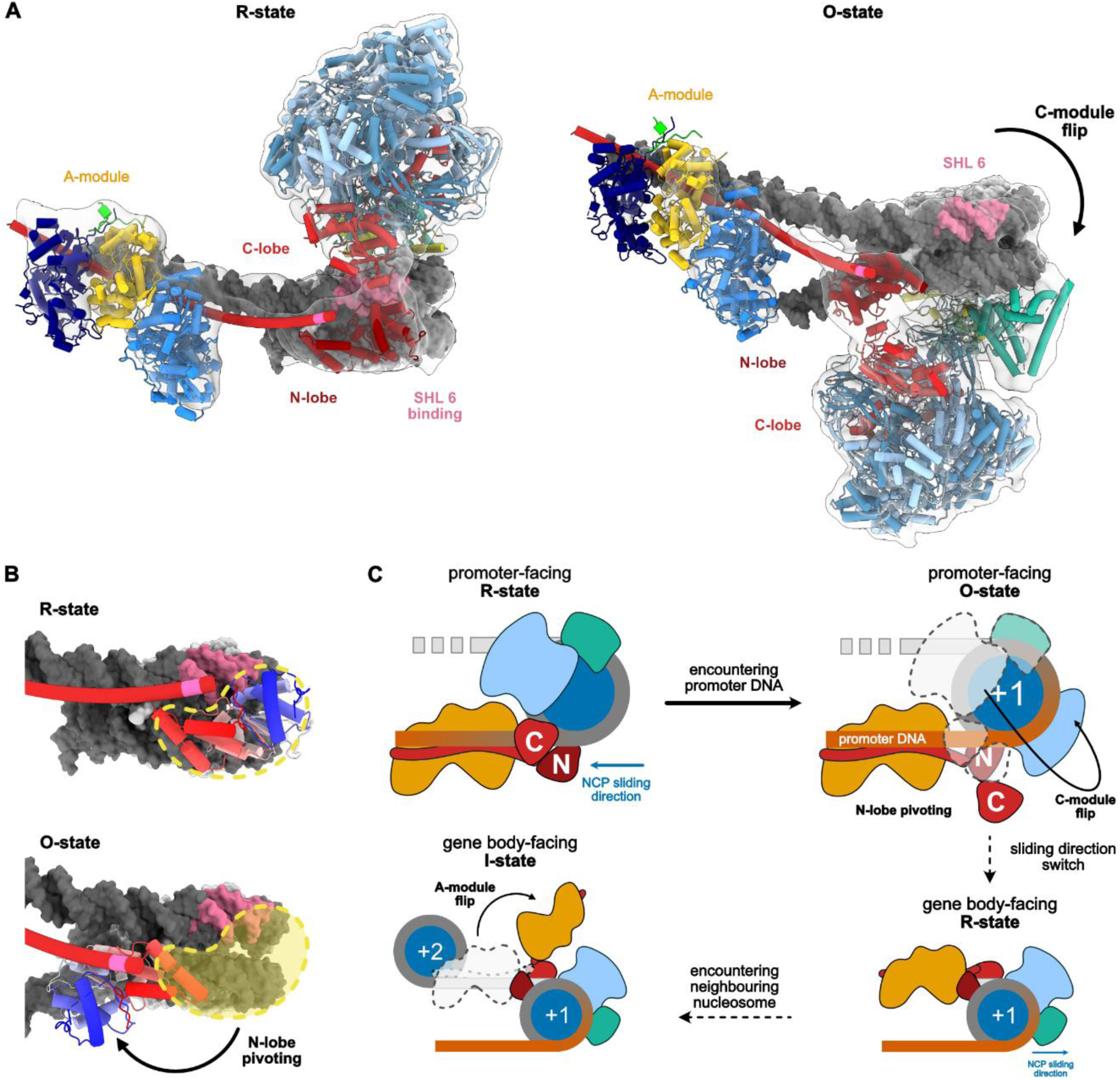
INO80 integrates nucleosomal and NDR sequence features into sliding arrest. **A,** Cryo-EM density map (EMD-xxx) with fitted model of R-state with an open ATPase (model is based on the AlphaFold 3 predictions and rigid-body fitted to the volume) and cryo-EM density map (low-pass filtered to 10 Å) with fitted model of the INO80 complex bound to the M2_rev_ nucleosome in O-state. Comparing R- to O-state shows that they are topologically related through a flip of the C-module on the nucleosome. The A-module can remain attached on extranucleosomal DNA, with the N-lobe of the Ino80 ATPase as pivot point. **B,** Depiction of N-lobe pivoting from R- to O-state. N-lobe location in R-state is marked yellow. **C,** Cartoon depiction of the interplay of all observed binding modes. Upon arrival at the +1 nucleosome site, intrinsic entry promoter DNA features (for example, A-tracts; marked dark orange) at the +1 site destabilize Ino80 ATPase engagement at SHL 6. This destabilization promotes flipping of the C-module into an inhibited O-state, while the A-module may remain attached to extranucleosomal DNA. Upon encountering the neighbouring nucleosome, shortening linker DNA sensing causes A-module flip and formation of an autoinhibited I-state, also restricting movement in an alternative way.

Comparison of the R- and O-states with A-modules engaged with extranucleosomal DNA now shows they are topologically related by a flip of the C-module across the nucleosome, rather than by differential binding of the A-module to extranucleosomal DNA. The N-lobe could act as the pivot point and change orientation between the nucleosomal gyres (Fig. 4 A, B; Movie S1). The C-module flip is facilitated by two alternative connections of the A-module to the N-lobe regulatory site via the post-HSA helix^33,47,48^, enabling the A-module to remain bound to extranucleosomal DNA in both R- and O-state. The compatibility with the O-state resolves the paradox of how A-module interaction with the NDR does not automatically lead to INO80 sliding the +1 nucleosome into the NDR.

## Discussion

Our results suggest a mechanism which explains how INO80 can position a +1 nucleosome by merely reading the DNA sequence and without the involvement of DNA-bound barrier factors. INO80 may stochastically slide a nucleosome from the gene body toward the unoccupied +1 nucleosome site, with the A-module engaging DNA ahead of the sliding direction. Upon arrival at the promoter, the R-state could be destabilized by DNA features, causing the INO80 C-module to flip into the sliding-incapable O-state and thereby arrest further upstream sliding of the +1 nucleosome. Our results are consistent with previous findings showing that, *in vivo*, in the absence of INO80, +1 nucleosome occupancy was observed further downstream in the gene body^20,49,50^.

Given sufficient intragenic linker DNA, the O-state may also serve as an intermediate toward a gene body-facing R-state. Such a reversal is plausible given INO80’s spin-rotating capacity^51,52^ and evidence that remodellers can swap nucleosome sides without fully detaching^53,54^. The combination of R-, O-, and I-states^33^ is well suited to explaining the formation of phased arrays, starting with the DNA-encoded kinetic trap at +1 and followed by subsequent spacing of +2, +3, and downstream nucleosomes (Fig. S8C). Here, the O- and I-states play distinct roles. The O-state could form in response to extra-/intranucleosomal DNA features, whereas the I-state could be induced in response to extranucleosomal DNA length, in particular sensing linker DNA shortening. A role of the A-module in setting the linker DNA length is consistent with altered linker lengths upon HSA helix mutations in the A-module^29^.

We and others previously noted that remodelling by INO80 depends on DNA sequence features, such as DNA shape, electrostatic potential, and rigidity^6,23,36,37,55^. This is also reflected, for example, in the composite propeller twist profile of all nucleosomes positioned by INO80 *in vitro* across the genome, which becomes asymmetric if sequences are sorted according to their shape profiles (Figs. 3F, S5-6). Intriguingly, the asymmetric propeller twist gradient in this profile is similar to that in the composite profile of all *in vivo* +1 nucleosome sequences sorted according to the direction of transcription^23^ (Fig. 3F), i.e., by a shape-independent sorting. Therefore, this similarity reflects the co-evolution of gene-oriented +1 nucleosome sequences and INO80’s sequence-dependent sliding directionality, so that INO80 does not slide nucleosomes into the promoter NDR but can generate +1 nucleosome positions, even on its own *in vitro*^6^. Our results now provide a mechanistic framework in which asymmetric nucleosomal DNA features can lead to orientation-dependent binding modes in R- and O-state, thereby driving directional sliding.

The *S. cerevisiae SWH1* +1 nucleosome enabled the discovery of this general mechanism, even though *SWH1* may represent an extreme case as a strict directional barrier for INO80, maybe because the propeller twist minimum is caused by an A-tract right at SHL 6/7 where the Ino80 ATPase needs to bind (Fig. S7). We found such sequence setup for some more *in vivo* +1 nucleosomes (Fig. S4), but skewed propeller twist is also generated by other sequences. More generally, nucleosomes positioned by INO80 *in vitro* or of other *in vivo* +1 nucleosomes (Fig. 3F) show a shallower slope of propeller twist and the propeller twist minimum is more upstream in the extranucleosomal linker/NDR. This may reflect that other sequences do not strictly stop sliding in one direction, but their similarly asymmetric DNA features still introduce a sliding direction bias, which amounts to a nucleosome positioning barrier^29^. In these cases, epigenetic marks, such as H3 tail modifications, and histone variants, together with barrier proteins, could provide additional layers of regulation and assist stable +1 nucleosome positioning *in vivo*.

In summary, our results provide a mechanism by which DNA features can directly regulate the binding mode of a core remodeller unit, leading to directional sliding and +1 nucleosome positioning. This provides a general model for how nucleosome organisation, with accurately positioned and phased nucleosomal arrays at promoters, can emerge from the readout of genome sequence information by remodellers.

## Data availability

For the cryo-EM structure and model of INO80 core bound to *SWH1* +1 NCP in R-state, the coordinates are deposited in the Protein Data Bank (PDB, https://www.wwpdb.org/) with the accession code xxxx; the cryo-EM density maps are deposited in the Electron Microscopy Data Bank (EMDB, https://www.ebi.ac.uk/emdb/) with the accession codes EMD-xxx (overall refinement), EMD-xxx (nucleosome refinement) and EMD-xxx (composite map). For INO80 core bound to array NCP in O-state in the ATP+AlF_x_ crosslinked dataset, the coordinates are deposited in the PDB with the accession code xxxx; the corresponding cryo-EM density maps are deposited in the EMDB with the accession codes EMD-xxx (overall refinement), EMD-xxx (nucleosome refinement) and EMD-xxx (composite map). For INO80 core bound to array NCP in R-state from this dataset, the accession codes are EMD-xxx (composite map), EMD-xxx (overall refinement) and EMD-xxx (nucleosome refinement).

For INO80 core bound to array NCP in I-state and R-state in the BeF_x_ dataset, the composite maps are deposited as EMD-xxx and EMD-xxx, respectively, along with EMD-xxx and EMD-xxx for overall refinements and EMD-xxx and EMD-xxx for nucleosome refinements. For INO80 core bound to *SWH1* +1 NCP in O-state, the accession codes are EMD-xxx (composite), EMD-xxx (overall refinement), and EMD-xxx (nucleosome refinement). The cryo-EM structure of INO80 core and A-module bound to M2_rev_ nucleosome in O-state is deposited with the accession code EMD-xxx.

All codes are available via our GitHub repository https://github.com/Hopfner-Lab/map2seq.

Plasmids and materials generated in this study are available upon request from the corresponding author.

## Acknowledgements

We are grateful to Manmohan Sharma and Cristian Rosales Hernández for processing advice, Felix J. Metzner and Vanessa Döbler for the preparation and initial processing of the M2_rev_ cryo-EM dataset, and Garp Linder and Ulrich Gerland for discussions. We thank Michael Kugler for technical support during cryo-EM data collection and Jan-Vincent Harre for support with GitHub depositions. M.L., A.B., and A.L.F. are grateful for the support from the International Max-Planck Research School for Molecules of Life (IMPRS-ML). D.S. is grateful for the support from the Ministry of Education and Science (MES) in Kosovo. We acknowledge Deutsche Forschungsgemeinschaft DFG (HO 2489/9-1, SFB1361-393547839, EXC 3113/1-533767322 to K.P.H. and SFB1064 to P.K. and K.P.H), the European Research Council (ERC Advanced Grant 833613 INO3D to K.P.H) and the National Institutes of Health, USA (R01GM108908 to B.B.).

## Author information

### Author notes

These authors contributed equally: Mariia Likhodeeva, Annika Brem.

### Contributions

M.L., A.B., A.L.-F., M.M., and O.F. conducted protein purification and DNA preparation. M.L., A.B., and A.L.-F. prepared cryo-EM samples. M.L., A.B., and A.L.-F. collected cryo-EM data with help from K.L.. M.L and A.B. conducted *in vitro* biochemistry assays. D.S. conducted genome-wide biochemistry assays, bioinformatics and derived DNA shape profiles under supervision of P.K.. M.L, A.B., A.L.-F., F.D., and K.-P.H. processed cryo-EM data with advice from K.L.. D.K. carried out structural model building and refinement. A.B. deposited the cryo-EM data. K.-P.H. wrote the scoring DNA sequences in cryo-EM maps method. M.L., A.B., and K.-P.H. analysed, interpreted and together with A.L.-F. and F.D. visualised the *in vitro* biochemical and structural data, and D.S. and P.K. visualised the shape analysis data. M.L. and K.-P.H. designed the studies. K.-P.H. supervised the overall project and together with P.K. and B.B. provided funding. M.L., A.B., and K.-P.H. wrote the initial draft and M.L., A.B, P.K., and K.-P.H. reviewed and edited the manuscript with contributions from all other authors.

### Declaration of generative AI and AI-assisted technologies in the writing process

Portions of this manuscript were edited to improve clarity and readibility using OpenAI’s ChatGPT. All such changes were proofread by the authors, scientific content and conclusions were made solely by the authors. The sequence-scoring Python code was written with the assistance of OpenAI’s ChatGPT.

## Materials and Methods

### Expression and purification of *S. cerevisiae* INO80^fl^, INO80^ΔN^ and INO80 A-module complexes from insect cells

Recombinant *S. cerevisiae* INO80^fl^ complex was generated as described previously^56^, using two pFBDM vectors. One vector contained the coding sequences of the INO80 C-module subunits (C-terminally 2xFLAG tagged Ino80, Rvb1, Rvb2, Ies6, Arp5) and a second vector contained the remaining coding sequences for the subunits of the A- and N-modules (Actin, Arp4, Arp8, Taf14, Ies2, Ies4, Ies1, Ies3, Ies5, and Nhp10). INO80^ΔN^, a complex in which truncation of the Ino80 N-terminus (amino acids 1-461) causes a loss of the Nhp10 module, was produced analogously using one vector containing the coding sequences of C-terminally 2xFLAG tagged Ino80(Δ1-461), Rvb1, Rvb2, Ies6, and Arp5 and a second vector containing the coding sequences for Actin, Arp4, Arp8, Taf14, Ies2, Ies4^57^. INO80 A-module, a complex in which C- and N-modules are truncated, was produced analogously using one vector containing the coding sequence of C-terminally 2xFlag tagged Ino80(330-598), and a second vector containing the coding sequences for Actin, Arp4, Arp8, Taf14, Ies2, Ies4^57^. Bacmids were generated using *E. coli* DH10 MultiBac cells. From each bacmid, baculoviruses were generated in *Spodoptera frugiperda* (Sf21) insect cells. Each baculovirus (1:200) was transferred to 1 l of *Trichoplusia ni* High Five culture for co-infection. Cells were cultured for 60 h at 27°C and harvested by centrifugation at 4°C.

For purification of the INO80^fl^ and INO80^ΔN^ complexes, cells were resuspended in lysis buffer containing 50 mM Tris HCl pH 8.0, 500 mM NaCl, 10% glycerol, 1 mM DTT, pepstatin A (0.28 μg/ml), PMSF (0.17 mg/ml), and benzamidine (0.33 mg/ml) and disrupted by sonication (3 × 1 min; duty cycle, 50%; and output control, 5 for INO80^fl^ and INO80^ΔN^. 4 x 45 s; duty cycle, 50%; and output control, 5 for INO80 A-module). The lysate was cleared by centrifugation at 34,500×g for 45 min at 4°C. The supernatant was incubated with 3 ml of ANTI-FLAG M2 Affinity Gel for 1 h and washed with 50 ml of Wash 1 buffer (25 mM HEPES pH 8.0, 500 mM KCl, 10% glycerol, 0.05% IGEPAL CA630, 4 mM MgCl_2_, and 0.25 mM DTT), 50 ml of Wash 2 buffer (25 mM HEPES pH 8.0, 200 mM KCl, 10% glycerol, 0.05% IGEPAL CA630, 4 mM MgCl_2_, and 0.25 mM DTT), and 10 ml of buffer A (25 mM HEPES pH 8.0, 150 mM KCl, 2 mM MgCl_2_, and 1 mM DTT). The protein was eluted from the matrix by incubation with 5 ml of buffer A supplemented with 0.2 mg/mL FLAG peptide in four incubation steps of 15 min each. The elution fractions were loaded onto a Mono Q 5/50 GL column and eluted by a linear salt gradient (150 mM KCl to 1 M KCl), resulting in highly pure INO80 complex. Aliquots were flash-frozen in liquid nitrogen in the presence of 20% of glycerol and stored at -80°C.

### Mononucleosome and nucleosomal array preparation

*H. sapiens* (*Hs*) histone H4 mutant H4K16Q were cloned into pET-21b vector and *X. laevis* (*Xl*) histones (H2A, H2B, H3, and H4) were cloned into pET-3a vector. Histones were purified as previously described^58^. In brief, histones were expressed in *E. coli* BL 21 (DE3) cells (Novagen) for 3 h at 37°C after induction at OD600 0.6 with 1 mM IPTG. Pelleted cells were disrupted by resuspension in wash buffer (50 mM Tris-HCl pH 8.0, 100 mM NaCl, 1 mM DTT) supplemented with 1 mg/ml lysozyme (Carl Roth, 8259), 1× protease inhibitor and 250 U benzonase (Merck Millipore, E1014) via three rounds of sonication. Inclusion bodies were washed two times with wash buffer supplemented with 1% Triton X-100 followed by two washes with standard wash buffer. Washed inclusion bodies were incubated with 1 ml DMSO for 30 min at room temperature followed by homogenization in resuspension buffer (7 M GdmCl, 20 mM sodium acetate pH 5.2, 1 mM EDTA) and 1 h incubation at room temperature. The supernatant was dialysed two times for 1.5 h against SAU 50 buffer (8 M urea; 20 mM sodium acetate, pH 5.2, 50 mM NaCl; 1 mM EDTA, 10 mM lysine). As first step of purification histones were separated by cation exchange chromatography (GE Healthcare HiTrap S HP) by applying a salt gradient. Histone-containing fractions were dialysed three times against refolding buffer (15 mM Tris-HCl, pH 8.0) of which one step needs to be over night to ensure proper refolding. Correct folded histones were finally separated by anion exchange chromatography (GE Healthcare HiTrap Q HP) using a salt gradient. Histone containing fractions were shock frozen in liquid nitrogen and lyophilized. WT *Hs*Histones were purchased from The Histone Source, Colorado State University.

Histones were resuspended in unfolding buffer (7 M guanidinium chloride, 25 mM Tris-HCl pH 7.5, 1 mM DTT). *Hs*H2A and *Hs*H2B were mixed in a 1:1 ratio for *Hs*H2A-H2B dimers; *Hs*H3.3 and *Hs*H4 were mixed in a 1:1 ratio for *Hs*H3.3-H4 tetramers (both for WT and H4 variants), and *Xl*H2A, *Xl*H2B, *Xl*H3 and *Xl*H4 were mixed in a 1.2:1.2:1:1 ratio for *Xl* octamers. Histones were dialysed against 2 changes of 1 L of refolding buffer (2 M NaCl, 25 mM Tris-HCl pH 7.5, 1 mM DTT) for 1.5 h at 4°C, then against 1 change of 1L of refolding buffer for 16 h at 4°C, and finally against 1 change of 1L of refolding buffer for 1.5 h at 4°C. Histone dimers, tetramers or octamers were purified by size exclusion chromatography using a Superdex 200 16/60 column (Cytiva). After concentrating in centrifugal filters (Amicon Ultra, 10 MW cutoff), histone dimers, tetramers or octamers were stored in 50% glycerol at -20°C.

DNA for nucleosome reconstitution was amplified by polymerase chain reaction (PCR) and purified by anion-exchange chromatography using a 1 mL HiTrap™ DEAE Sepharose™ Fast Flow column (Cytiva). The DNA was subsequently precipitated by ethanol precipitation and resuspended in H_2_O.

For reconstituting nucleosomes, DNA was mixed at a 1.1-fold molar excess with the histone *Xl* octamer (80S80 nucleosomes) and with *Hs* dimer and *Hs* tetramer (for the rest) in a buffer containing 25 mM HEPES pH 7.5, 0.25 mM DTT, and 2 M NaCl. The NaCl concentration was gradually decreased to 50 mM over 16 h at 4°C by salt gradient dialysis. After this, nucleosomes were purified by anion exchange chromatography using a 1 ml Resource Q column, and fractions containing nucleosomes were pooled and dialysed to 25 mM Tris-HCl pH 7.5, 0.25 mM DTT and 50 mM NaCl, concentrated to 1 mg/ml (Amicon Ultra, 30 MW cutoff), and stored at 4°C or aliquoted and flash-frozen in liquid nitrogen in the presence of 10% of glycerol and stored at -80°C.

DNA for array reconstitution was amplified by polymerase chain reaction (PCR) and purified using NucleoSpin Gel and PCR Clean-up, Maxi kit for gel extraction and PCR clean up (Macherey-Nagel, 740610.20).

For reconstituting arrays, DNA was mixed at a 0.83-fold molar excess with the histone dimer and tetramer in presence of recombinant albumin in a HEN2000 buffer containing 10 mM HEPES, pH 8; 0.1 mM EDTA; 2 M NaCl. The NaCl concentration was gradually decreased to 50 mM over 16 h at 4°C by salt gradient dialysis. After this, arrays were dialysed to HEN50 buffer (10 mM HEPES, pH 8; 0.1 mM EDTA; 50 mM NaCl), concentrated using Amicon Ultra, 30 MW cutoff, and stored at 4°C.

### MNase array quality control

Correct nucleosomal array reconstitution was assessed by performing a MNase digest. Here, 1000 ng of array was digested with 20 U MNase (NEB, M0247S) using provided reaction buffer. The reaction was stopped using STOP-buffer (50 mM Tris pH 7.5, 4% SDS, 100 mM EDTA pH 8.0) at different incubation time points (1’ and 5’). Afterward samples were incubated with Proteinase K (Thermo Fisher Scientific, EO0491) for 1 h at 50°C and digested DNA was isolated using ethanol precipitation in the presence of glycogen (Roche, 10901393001). The isolated DNA fragments were separated using 1.5% (Biozym, 840004) agarose gel electrophoresis with 1×TAE (40 mM Tris pH 8, 1 mM EDTA, 20 mM acetic acid) as running buffer (120 V, 1.5 h) and stained with GelRed (Biotium, 41003). Gels were imaged using a GE Healthcare Typhoon FLA9000 imager.

### Restriction enzyme accessibility assays

50 nM of INO80^ΔΝ^ and 12.5 nM arrays (assembled with WT *Hs* octamer) were mixed in a reaction buffer containing 40 mM Tris pH 7.5, 60 mM KCl, 5 mM MgCl_2_, and 2 U/μL EcoRV. Each mixture was divided in two after a 2-minute preincubation at 30°C: the control without ATP and the reaction that was started with the addition of 1 mM ATP. Samples were quenched at various time-points with equal volumes of 20 mM Tris pH 7.5, 2% SDS, 70 mM EDTA and 20% glycerol and incubated with Proteinase K for 30 minutes at 50°C to digest all proteins. The DNA fragments were then isolated by ethanol precipitation using glycogen (Roche, 10901393001). The isolated DNA (cut and uncut fragments) for each timepoint were resolved using 1.5% (Biozym, 840004) agarose gel electrophoresis with 1×TAE as running buffer (120 V, 1.5 h) and stained with GelRed (Biotium, 41003). Gels were imaged using a GE Healthcare Typhoon FLA9000 imager and were evaluated using ImageJ. After background subtraction, the total intensities of the cut and uncut band peaks were evaluated and the percentages of the fraction cut for each sample were plotted with Prism (GraphPad Software).

### NADH-coupled ATPase assay

The ATP hydrolysis rate of INO80^ΔN^ was determined using an NADH-coupled ATPase assay. Reactions were carried out in a total volume of 50 μL containing 30 nM INO80^ΔN^ in assay buffer composed of 25 mM HEPES (pH 8.0), 50 mM KCl, 1 mM DTT, 2 mM MgCl₂, and 0.1 mg/mL BSA. The reaction mixture was supplemented with 0.5 mM phosphoenolpyruvate, 1 mM ATP, 0.1 mM NADH, 25 U/mL lactate dehydrogenase, and pyruvate kinase. Assays were performed at 25°C in non-binding black 384-well plates, and NADH consumption was monitored fluorometrically over one hour using a Tecan Infinite M100 plate reader (excitation 340 nm, emission 460 nm). Where indicated, ATPase activity was measured in the presence of 200 nM nucleosome or 50 nM of tetranucleosomal array. All reactions were performed in triplicate. ATP turnover rates were determined from the maximal initial linear rates of the NADH depletion curves and corrected by subtraction of a buffer blank.

### Nucleosome sliding assays

Nucleosomes with 6’FAM–labelled extranucleosomal DNA were used for monitoring the sliding activity of purified recombinant INO80^ΔN^ complex. Nucleosome (150 nM) was incubated with 50 nM INO80^ΔN^ complex in sliding buffer [25 mM HEPES pH 8.0, 60 mM KCl, 7% glycerol, recombinant albumin (0.10 mg/ml), 0.25 mM DTT] at 25°C. The sliding reaction was started with the addition of 1 mM ATP and 2 mM MgCl_2_ and stopped at indicated time points by addition of Lambda DNA (0.2 mg/ml). Nucleosome species were separated by native polyacrylamide gel electrophoresis (PAGE) on a 3 to 12% acrylamide Bis-Tris gel (Invitrogen) and visualised using the Typhoon imaging system (GE Healthcare). Experiments were performed in triplicate.

Nucleosome sliding assays were quantified using ImageJ by measuring band intensities corresponding to unremodelled nucleosomes (end-positioned; lowest band), sliding intermediates, and fully remodelled nucleosomes (centrally positioned; uppermost band). The fraction of centrally positioned nucleosomes was calculated as the intensity of the fully remodelled species expressed as a percentage of the total signal. The remodelled fraction was defined as the combined intensity of sliding intermediates and fully remodelled nucleosomes relative to the total intensity. Kinetic data of the remodelled fraction were fitted using a one-phase exponential decay model: *y*(*t*) = (*y*_0_ − *p*)*e*^−*Kt*^ + *p*, the centrally positioned nucleosome data was fitted with *y*(*t*) = *y_inf_*(1 − *e*^−*Kt*^) for visualisation purposes.

### Fluorescence anisotropy

Increasing concentrations of INO80 A-module (0, 8, 16, 32, 64, 128, 256 and 512 nM final) were prepared in assay buffer (25 mM HEPES, pH 8.0, 120 mM KCl, 2 mM MgCl₂, 2% glycerol, 0.01% Triton X-100, 1 mM DTT) and mixed at a 1:1 (v/v) ratio with 50-bp 6-FAM-labelled DNA (50P or 50G; Table Sxxx) to yield a final DNA concentration of 5 nM in a total volume of 20 μl (Greiner flat-bottom black 384-well plates). Samples were incubated for 20 min at room temperature prior to measurement of fluorescence anisotropy (λ_ex_ = 470 nm, λ_em_ = 520 nm) using a TECAN Infinite M1000 plate reader. All measurements were performed in triplicate.

For data analysis, background signal from the 0 nM protein control was subtracted from all data points. The resulting datasets were analysed using Prism (GraphPad Software) and fitted with a non-linear, non-cooperative 1:1 binding model 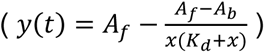, where *y* denotes anisotropy, *A_f_* and *A_b_* represent the anisotropy of free and bound ligand, respectively, *K_d_* is the dissociation constant, and *x* corresponds to receptor concentration. Apparent dissociation constants were derived from the fitted parameters.

### Vitrification for cryo-EM

For INO80^ΔN^-array NCP complexes, 600 nM INO80 and 150 nM of arrays were mixed in cryo-EM buffer (20 mM HEPES, pH 8.0; 60 mM KCl; 2 mM MgCl_2_; and 1 mM DTT). 1 mM ATP was added to the mixture and the sample was incubated at 25 degrees for 10 min, then 1.5 mM of the respective nucleotide (AlF_x_ or BeF_x_) was added. For INO80^fl^-80N0 M2_rev_ nucleosome, INO80^fl^-80S80 nucleosome and INO80^ΔN^-80S0 nucleosome complexes; 500 nM INO80 and 500 nM nucleosome, 150 nM INO80 and 165 nM nucleosome, 600 nM INO80 and 600 nM nucleosome respectively were mixed in cryo-EM buffer. Then 1 mM ADP-AlF_x_ (M2_rev_, 80S80) or 1.5 mM ADP-AlF_x_ (80S0) was added.

After the addition of the respective nucleotide, samples were incubated at 25 degrees for 10 min. In case of crosslinking (for AlF_x_ XL dataset), 0.1% glutaraldehyde was added and sample was incubated on ice for 10 min. Octyl-β-glucoside was added (0.045%), and 4.5 μl was applied onto a glow discharged Quantifoil R2/1 + 2 nm C Cu200 grid (arrays, 80S80, 80S0) or onto a glow discharged Quantifoil R2/1 Cu200 grid (M2_rev_). The sample was vitrified in liquid ethane using an EM GP plunge freezer (Leica; 10°C and 90% humidity; 2.2 s blot; carbon grids were pre-blotted for 15 s).

### Cryo-EM data collection

For INO80^ΔN^-array NCP, INO80^fl^-80S80 nucleosome and INO80^ΔN^-80S0 nucleosome complexes, movies of particles embedded in vitreous ice were collected at liquid nitrogen temperature using a FEI Titan Krios G3 transmission electron microscope (300 kV, Thermo Fisher Scientific) equipped with a Selectris X imagining filter (slit width 10 eV) with a Falcon4 direct electron detector. The movies were recorded in counting mode using EPU acquisition software (Thermo Fisher Scientific) at ×165,000 magnification with a pixel size of 0.727 Å/pixel and nominal defocus range of -0.5 to -2.6 μm. The total electron dosage of each movie was 40 e/Å^2^, fractionated into 40 movie frames.

For INO80^fl^-80N0 M2_rev_ nucleosome complexes, movies of particles embedded in vitreous ice were collected at liquid nitrogen temperature using an FEI Titan Krios G3 transmission electron microscope (300 kV) equipped with a GIF quantum energy filter (slit width 20 eV) and a Gatan K2 Summit direct electron detector. The movies were recorded in counting mode using EPU acquisition software (Thermo Fisher Scientific) at ×130,000 magnification with a pixel size of 1.059 Å/pixel and nominal defocus range of -1.1 to -2.9 μm. The total electron dosage of each movie was 41.04 e/Å^2^, fractionated into 40 movie frames.

### Cryo-EM data processing

For all datasets, movie frames were motion-corrected using MotionCor2^59^. All subsequent processing steps were performed in cryoSPARC (v5.0.6 and former versions)^60^, and the resolutions reported here are calculated based on the gold-standard Fourier shell correlation criterion (FSC = 0.143) with 3DFSC plots using Remote 3DFSC Processing Server^61^ or with cryoSPARC^60^. CTF parameters of each dataset were determined using patch CTF estimation (multi) in cryoSPARC^60^. The data collection and refinement statistics for each dataset are summarized in the cryo-EM validation tables 2-5. Figures and composite maps were generated in UCSF ChimeraX 1.5 or 1.6.145.32^62^.

For the INO80^ΔN^-array complex AlF_x_ XL dataset, 40195 micrographs were collected. The processing scheme is depicted in Fig. S9. Initial particle picking was done using Blob picker (2621k particles). Particles were subjected to 2D classification, ab initio reconstruction, and heterogeneous refinement (1069k particles). Particles of INO80-NCP complexes, were then extracted with a box size of 600 px and a pixel size of 0.727 Å/pixel, and used for 3D-classification, sorting the particles in O-(259k particles) and R-states (134k particles). Particles of both states were further 3D-classified and particles with clearly defined features were used for homogeneous and/or non-uniform refinement of R-state^63^. The final resolution for INO80-NCP complex in O-state overall and local refinements was 2.91 and 3.49 Å, respectively^61^. The final resolution for INO80-NCP complex in R-state overall and local refinements was 2.68 and 2.98 Å, respectively^61^.

For the INO80^ΔN^-array complex BeF_x_ dataset, 31372 micrographs were collected. The processing scheme is depicted in Fig. S10. Initial particle picking was done using Blob picker (1332k particles). Particles were subjected to 2D classification, ab initio reconstruction, and heterogeneous refinement (198k particles). Particles of INO80-NCP complexes were used for template picking^64^, 2D-, and 3D-classification, ab initio reconstruction and heterogeneous refinement. Particles were extracted with a box size of 600 px and a pixel size of 0.727 Å/pixel. 708k particles were used for homogeneous refinement of R-state. The final resolution overall and local refinements for INO80-NCP in the R-state was 2.52 and 2.43 Å, respectively^60^. Particles of INO80-NCP in the R-state were further 3D-classified and particles of classes with the A-module density were then re-extraction with a box size of 800 px and refined. The final resolution for INO80-NCP complex in the I-state both overall and local refinements was 3 Å^60^.

For the INO80^ΔN^-80S0 dataset, 16603 micrographs were collected. The processing scheme is depicted in Fig. S11. Initial particle picking was done using Blob picker (442k particles). Particles were subjected to 2D classification, ab initio reconstruction, and heterogeneous refinement (198k particles). Further 2D-, and 3D-classification, and homogeneous refinement were applied to sort the particles leading to the sharpest reconstruction. 150k particles were then extracted with a box size of 600 px and a pixel size of 0.727 Å/pixel and used for homogeneous refinement of O-state. The final resolution for INO80^ΔN^-80S0 complex in the O-state overall and local refinements was 2.68 and 3.25 Å, respectively^60^.

For the INO80^fl^-80S80 nucleosome dataset 12000 micrographs were collected. Initial particle picking was done using Blob picker (9249k particles). For INO80^fl^-80S80 nucleosome complex (the processing scheme is depicted in Fig. S12), particles were subjected to 2D classification, ab initio reconstruction, and heterogeneous refinement (226k particles). Particles with clearly defined features, were used for as input for a Topaz train job^65^. After four rounds of Topaz, particles were extracted with a box size of 504 px and a pixel size of 0.727 Å/pixel. The particles were subjected to 2D classification and heterogeneous refinement. The class that showed the most defined features was selected (172k particles) and used for ab-initio reconstruction and non-uniform refinement^63^ and further heterogeneous refinement into three classes. One class with 127k particles was used for homogeneous and local refinements. The final resolution of the last homogeneous refinement and the last local refinements were 2.89 and 2.75 Å, respectively^61^.

For 80S80 nucleosome (the processing scheme is depicted in Fig. S13), particles were subjected to 2D classification, and particles with clearly defined features, were used for as input for a Topaz train job^65^. After four rounds of Topaz, particles were extracted with a box size of 360 px and a pixel size of 0.727 Å/pixel. The particles were subjected to 2D classification and heterogeneous refinement. The class that showed the most defined features was selected (650k particles) and used for further heterogeneous refinement into four classes. One class with 450 classes was used for non-uniform refinement^63^, 3D classification, and a extra round of heterogeneous refinement. Three classes with 82k, 121k and 112k particles were selected and used for non-uniform refinement^63^, first separately and then combined. The combined particles were used for 1 round of 3D classification with 2 classes. One class with 205k particles was selected and used for local refinement. The final resolution of the last local refinement was 2.81 Å ^61^.

For the INO80^fl^-80N0 M2_rev_ nucleosome dataset, 14722 micrographs were collected. The processing scheme is depicted in Fig. S14. Initial particle picking was done using Blob picker (4739k particles). Particles were subjected to 2D classification, ab initio reconstruction, and heterogeneous refinement (355k particles). Particles with clearly defined features, were used for two rounds of Topaz^65^, 2D-, and 3D-classification, ab initio reconstruction and heterogeneous refinement. 55k particles were used for non-uniform refinement^63^ and further 2D classification and ab initio reconstructions, followed by heterogeneous refinement into two classes. One class with 25k particles was used for 3D variability analysis. 14k particles were used for non-uniform and local refinements^63^, 3D flexibility analysis^66^ and final non-uniform refinement with a box size of 512 px and a pixel size of 1.059 Å/pixel. The final resolution of the INO80^fl^-80N0 M2_rev_ nucleosome complex was 4.37 Å ^61^.

### Model building and refinement

Model templates were taken from AlphaFold2 34265844 model predictions for all *S. cerevisiae* protein chains, and from the cryo-EM INO80 Core structure for the *H. sapiens* histones (6FML)^67^, and from the cryo-EM Nucleosome structure for the *X. laevis* histones (6WZ5)^68^, respectively. The DNA model template was also taken from the cryo-EM Nucleosome structure for *X. laevis* histones (6WZ5)^68^ and mutated to the *S. cerevisiae* natural promoter sequence as described in the Suppl. Method "Mapping Nucleosomal DNA".

All model templates were manually placed into the corresponding composite cryo-EM maps, followed by rigid-body refinement with ChimeraX^62^. The initial models were then energy-minimised using ISOLDE^69^ and manually inspected and corrected with Coot^70^.

Reciprocal space refinement was done at 2.8 Å resolution using jelly-body restraints with Servalcat^71^ against the calculated structure factors from the composite cryo-EM maps, resulting in R=0.254, <FSC>=0.871, rmsd bonds=0.012 Å, rmsd angles=1.61° for the INO80 core bound to array NCP in O-state, and in R=0.262, <FSC>=0.844, rmsd bonds=0.011 Å, rmsd angles=1.68° for the INO80 core bound to 80S80 *SWH1* +1 NCP in R-state.

### Genome-wide chromatin remodelling assay

*S. cerevisiae* genomic plasmid library^72^ chromatin was reconstituted with embryonic *D. melanogaster* histones by salt-gradient dialysis (SGD) under low assembly degree conditions and remodelled by INO80 as described^56^ with the following modifications. DTT (2 mM) was used as reducing agent instead of β-mercaptoethanol during SGD. Remodelling reactions were done with 600 ng whole-genome plasmid library DNA reconstituted as SGD, 20 nM recombinant *S. cerevisiae* WT INO80^fl^, 200 U KpnI (only replicates 1 and 2, Fig. S6) in a total volume of 50 µl for 2 h in the presence of ATP. For replicates 2 and 3, the remodelling reaction buffer did not contain ammonium sulphate.

### DNA shape analysis

Genome-wide nucleosome dyad positions were called from paired-end Illumina (Next Seq 1000 for replicates 1 and 2, NextSeq 2000 for replicate 3) MNase-seq data. MNase-seq read pairs were mapped using Bowtie2 (2.5.4) to the yeast reference genome (SacCer3, R64-1-1), filtered for nucleosomal size fragments (130 to 180 bp) and trimmed to 50 bp around the inferred dyad (calculated from paired-end fragment midpoints). Read counts were normalized by subsampling each group of samples to the number of reads in the sample with the lowest sequencing depth (target_n).

Nucleosome peak positions in the aggregated MNase-seq data were identified on dyad-centred coverages using the NucleR (2.42.0) peak-calling algorithm (Bioconductor). Only the top 3% of dyad positions ranked by peak height (97% threshold) were used as high-confidence nucleosome dyads. These positions were stored as .rds or .bed files for downstream analysis.

A matrix of DNA shape features was computed around each dyad using the DNAshapeR (1.38.0) package within a symmetrical window of ±160 bp (halfwin = 160L). For direct comparison with^56^, we focused on the propeller twist (ProT) feature and generated dyad-aligned composite plots. These composite plots gave approximately symmetrical profiles. We reasoned that the symmetry could stem from the overlay of randomly oriented asymmetric profiles and sought to disentangle sequence orientations by DNA shape-based clustering (Fig. S5). Each dyad-centred ProT profile was Z-score-transformed row-wise such that each dyad profile had mean 0 and standard deviation 1. Only the profiles were retained where at least 95% of positions had finite Z-scores. The ProT matrix was used to compute a Euclidean distance matrix, which was clustered via hierarchical clustering with Ward.D2 linkage yielding a dendrogram based on ProT patterns of nucleosomal sequences. At the level of k = 2 clusters, we got two distinct clusters of nucleosomes. The right-side-heavy cluster, which showed the lowest ProT in the 0 to +160 bp region, appeared as mirror image of the left-side-heavy cluster profile and was flipped and merged with the left-side-heavy cluster to generate the final composite profile (plots were smoothed with 41 bp rolling mean for visualization – zoo/rollmean (1.8.14)). The ProT profile was also calculated for the sequence of the *SWH1* in vivo +1 nucleosome dyad ±160 bp.

## Supplementary Information

**Supplementary Fig. 1:**
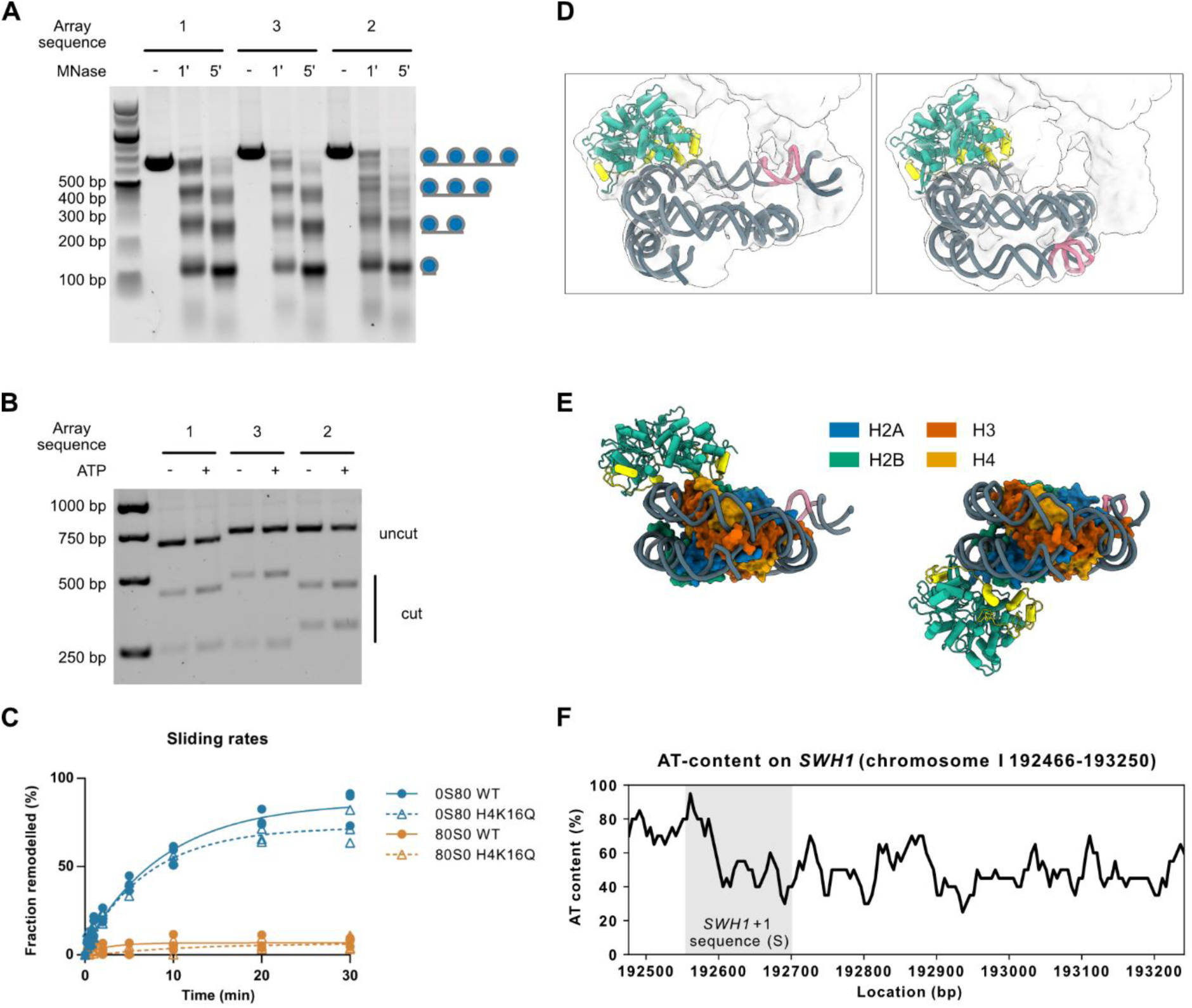
Supplementary array data. **A**, Agarose gel of micrococcal nuclease (MNase)-digested nucleosomal arrays assembled on natural sequences, indicating a regular nucleosomal structure. **B**, Representative agarose gel of EcoRV restriction enzyme accessibility assay. **C**, Comparison between H4K16Q and WT octamer 0S80 and 80S0 nucleosomes sliding. Arp5-Ies6 grip in R- and O-states shows similar engagement with DNA (**D**) and different engagement with histones (**E**). **F**, AT-content of the natural sequences chosen for the array assembly.

**Supplementary Fig. 2:**
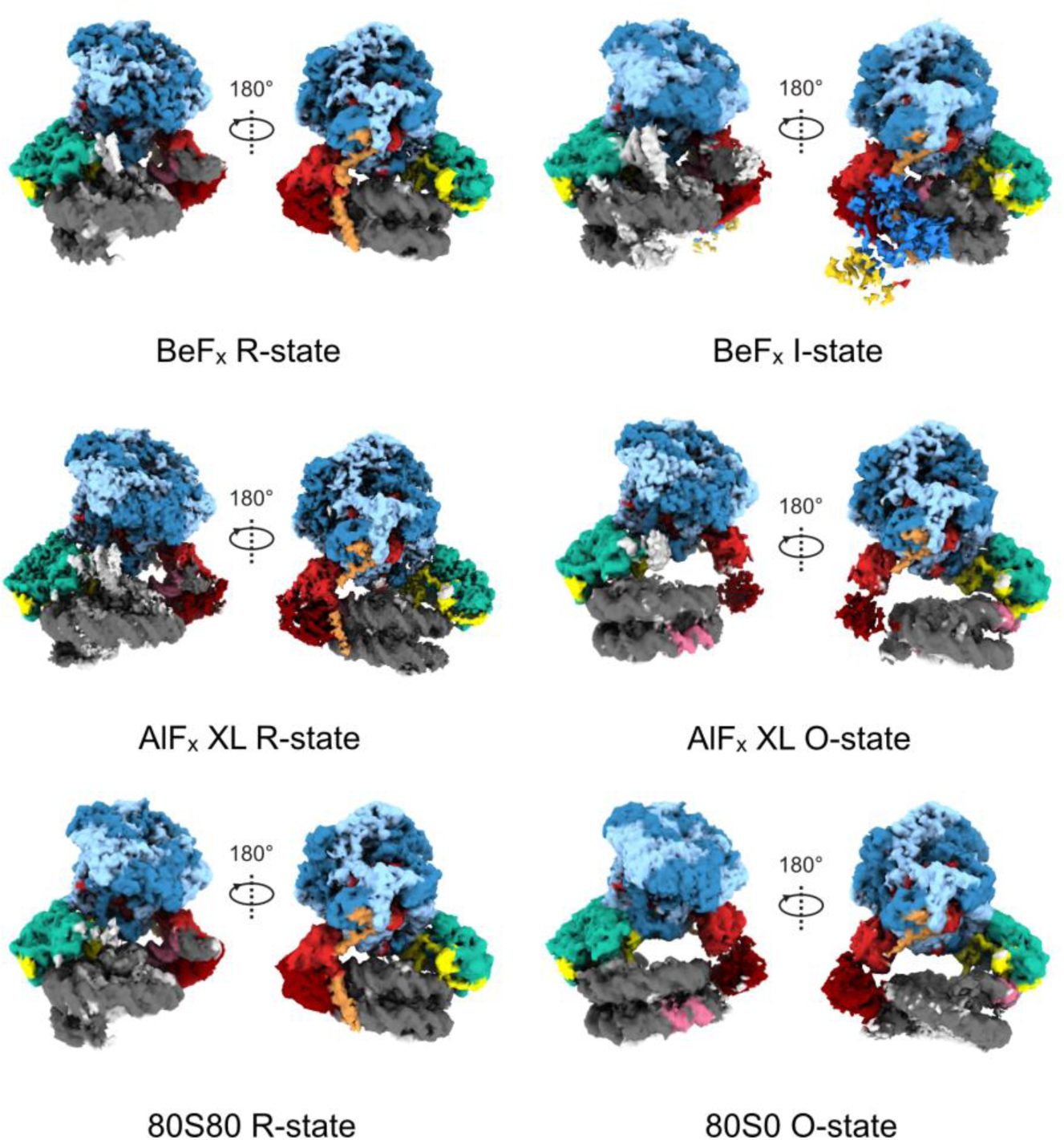
Cryo-EM volumes obtained of INO80 on arrays and *SWH1* +1 nucleosomes. Maps are coloured according to their models.

**Supplementary Fig. 3:**
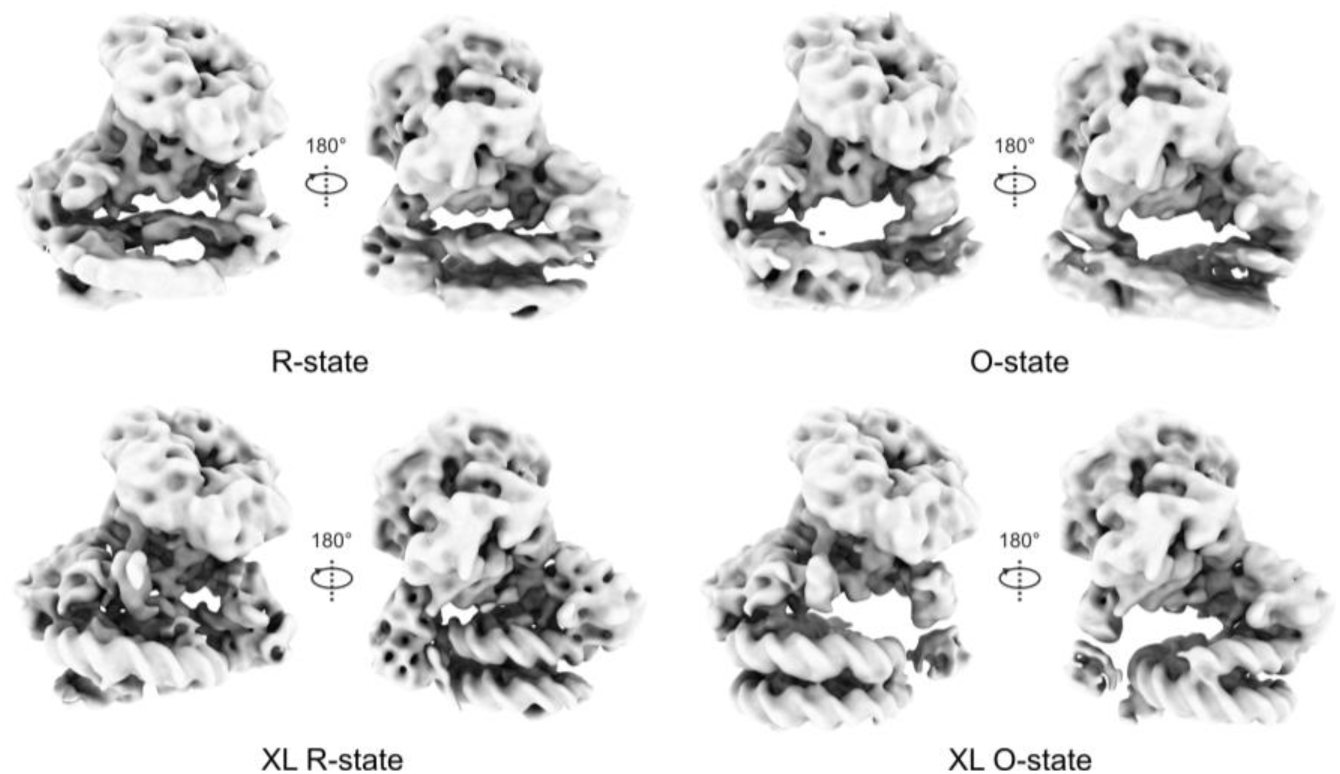
Cryo-EM density maps (low-pass filtered to 10 Å) of R- and O-states in the AlF_x_ datasets without and with glutaraldehyde present. The O-state is present in the AlF_x_ dataset, but it is less stable, as its presence could be increased by mild glutaraldehyde crosslinking. XL, crosslinked.

**Supplementary Fig. 4:**
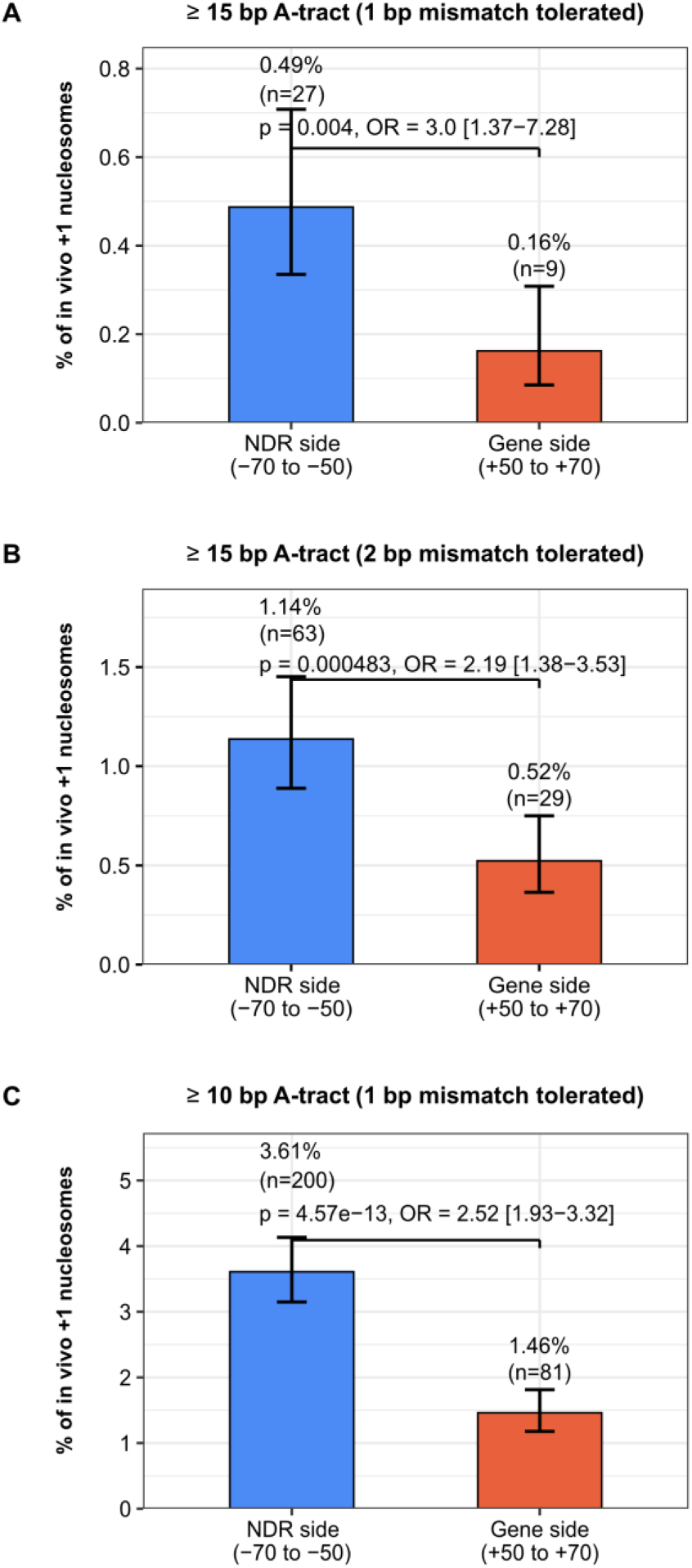
Enrichment of long A-tracts at up- vs. downstream flank in +1 nucleosomes *in vivo*. Long A-tracts were identified in the *S. cerevisiae* genome under three criteria: **A,** ≥15 consecutive adenines with 1 bp mismatch tolerance, **B,** ≥15 consecutive adenines with 2 bp mismatch tolerance, and **C,** ≥10 consecutive adenines with 1 bp mismatch tolerance. For each criterion, the occurrence of A-tracts was assessed relative to the +1-nucleosome dyad either on the NFR side (−50 to −70 bp) or on the gene side (+50 to +70 bp). Coordinates for +1 nucleosome positions were based on Chereji et al.^1^. Enrichment was tested by Fisher’s exact test; odds ratios (OR) and p-values are reported.

**Supplementary Fig. 5:**
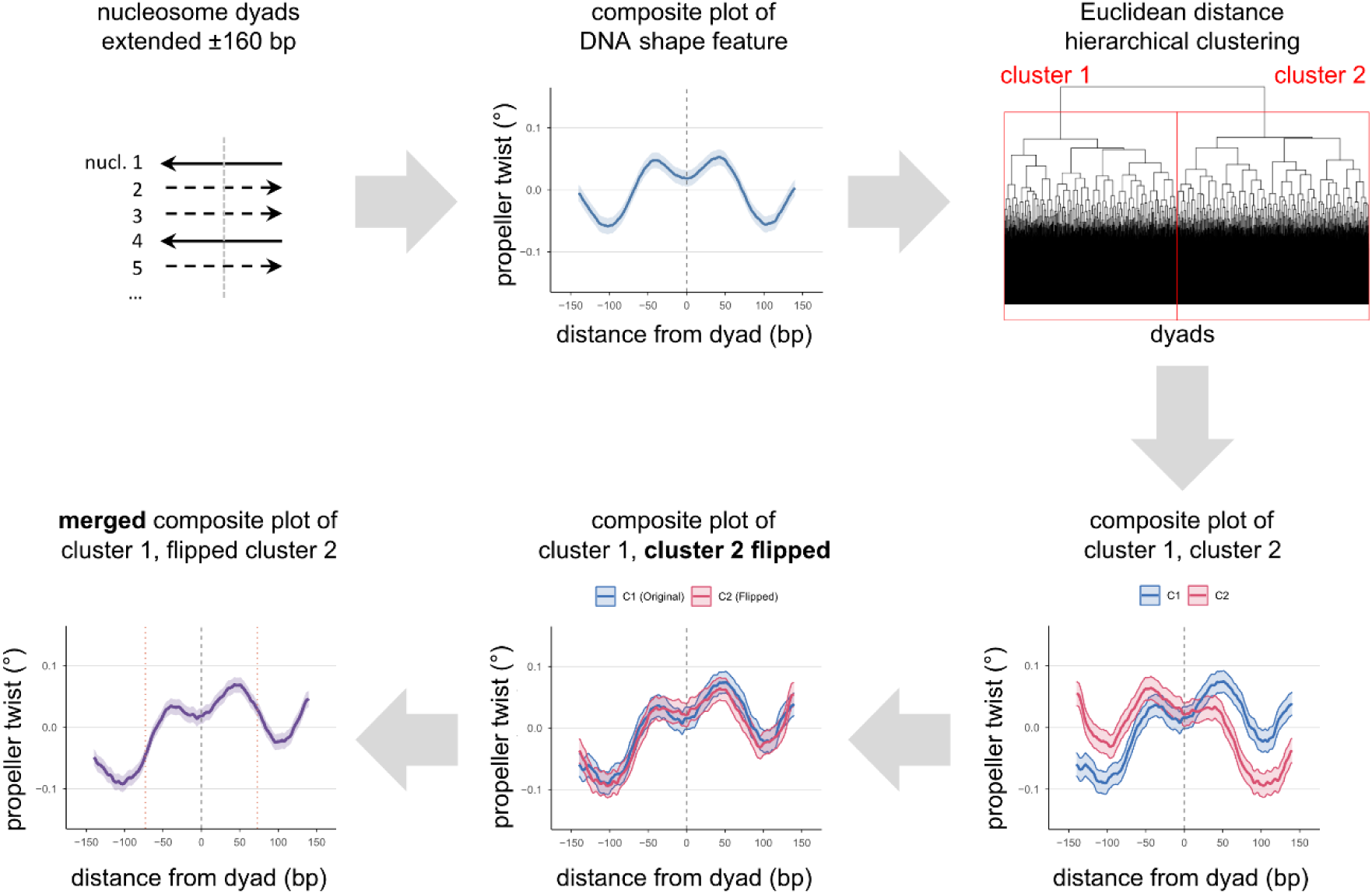
Orientation of DNA shape profiles based on Euclidean distance hierarchical clustering. DNA shape profiles were computed for extended DNA sequences aligned at nucleosome dyads called from MNase-seq data and plotted as dyad-aligned composite plot, which yields at first a rather symmetrical composite profile (Z-scored, smoothed). Hierarchical clustering based on Euclidean distances was computed for individual DNA shape profiles and plotted as dendrogram. Composite profiles of the two main clusters 1 and 2 (C1, C2) appeared like mirror images reflected at the dyad axis. Flipping the composite profile of C2 around the dyad axis and merging it with the composite profile of C1 yielded the composite profile of sequences in all nucleosomes positioned by INO80 *in vitro* and oriented by DNA shape feature (Fig. 3F).

**Supplementary Fig. 6:**
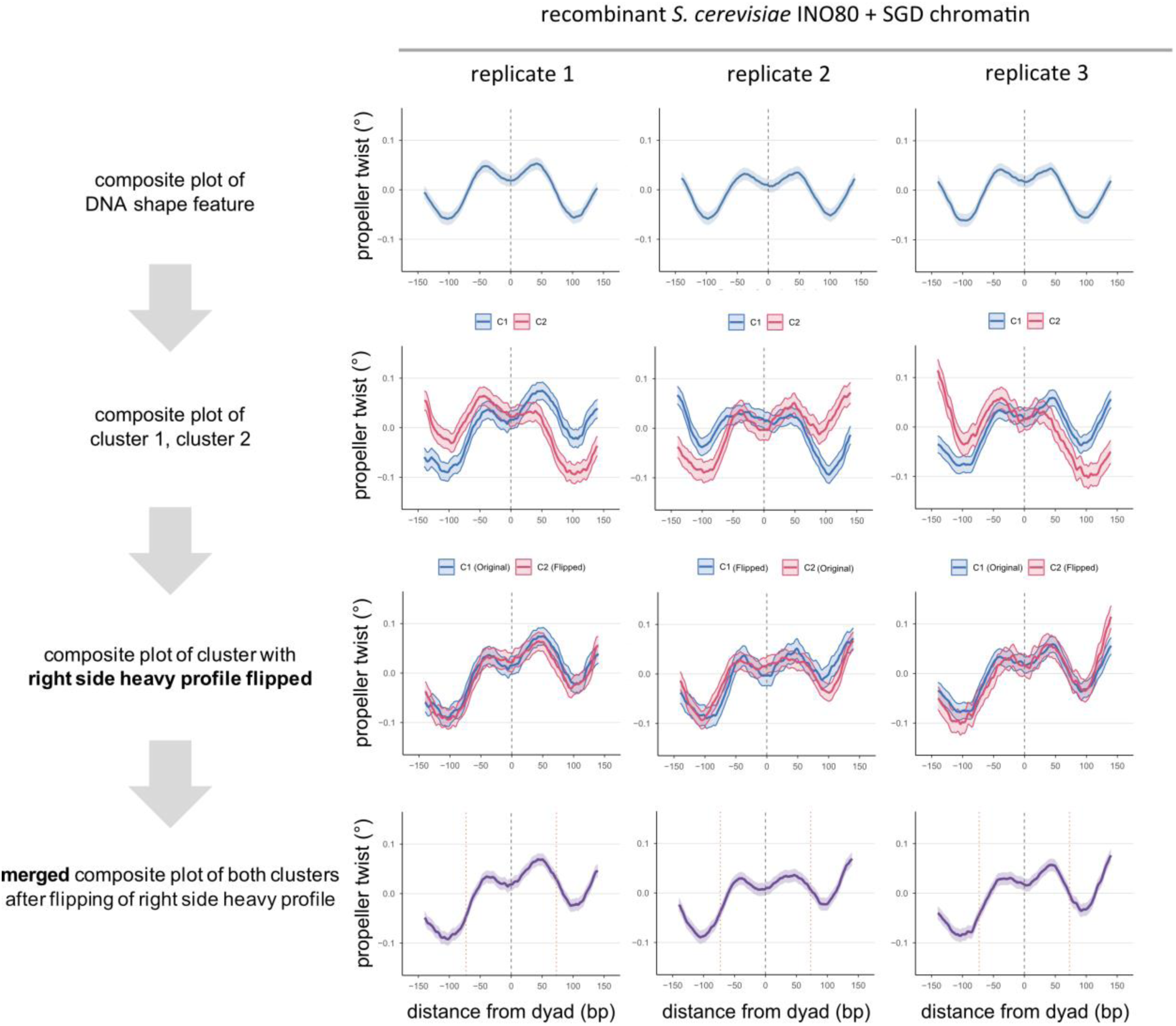
Replicates of DNA shape-oriented DNA shape profiles for nucleosomes positioned *in vitro* by INO80. As Fig. S5 for three replicates. Replicate 1 is the same as in Fig. S5. Replicate 2 was generated by an independent remodelling reaction from the same SGD chromatin as replicate 1. Replicate 3 was generated by a remodelling reaction from an independent SGD chromatin reconstitution.

**Supplementary Fig. 7:**
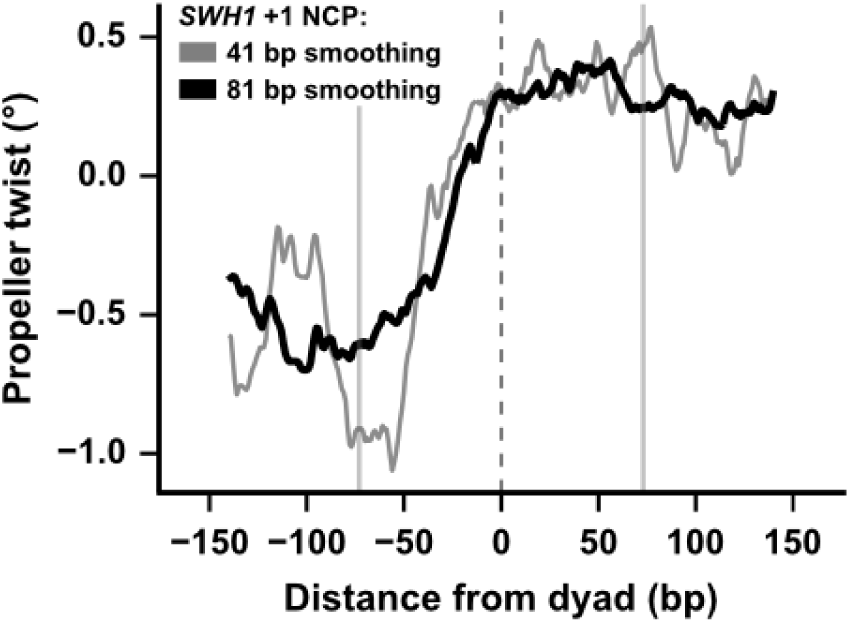
*SWH1* +1 nucleosome also shows an asymmetric skew of propeller twist similar to all *in vivo* +1 nucleosomes and all nucleosomes positioned by INO80 *in vitro* (Fig. 3F). DNA shape (propeller twist, Z-score) profile for the sequence centred at the dyad of the *in vivo SWH1* gene. For comparison of this profile for an individual sequence with the composite profiles for many sequences (all *in vivo* +1 nucleosomes and all nucleosomes positioned by INO80 *in vitro*, Fig. 3F), both the same (grey trace, 41 bp rolling mean) and increased (black trace, 81 bp rolling mean) smoothing is shown. Increased smoothing mimics the added effective smoothing in the composite plots due to averaging over many sequences (Fig. 3F). Vertical dashed line marks the dyad, vertical solid lines the nucleosome borders (+/- 73 bp from dyad).

**Supplementary Fig. 8:**
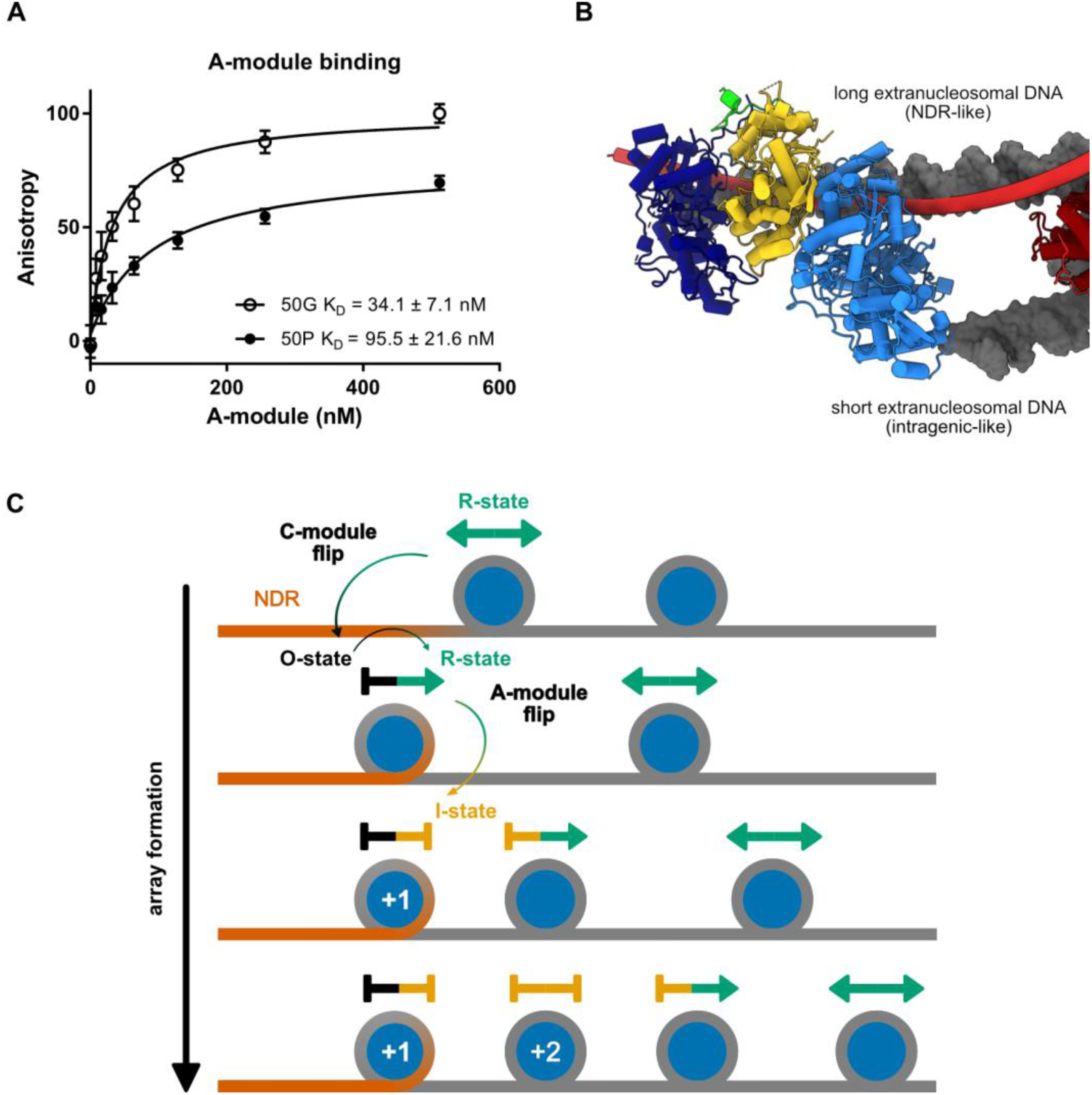
INO80 +1 nucleosome positioning mechanism. **A,** Fluorescence anisotropy of A-module binding to 50 bp of either promoter-(50P) or gene body-facing (50G) DNA (see Fig. 3B). **B,** Arp8 of the A-module engages both extranucleosomal DNA strands in O-state binding to the M2_rev_. The long extranucleosomal DNA (equivalent to the “NDR” in the *SWH1* studies) was bound along the Ino80 HSA domain and Arp8 N-terminus. An unexpected second DNA interaction site on Arp8 bound the short extranucleosomal DNA (equivalent to intragenic linker in the *SWH1* studies), which helped stabilize the overall arrangement to an extent that it could be visualized by cryo-EM. **C,** INO80 may stochastically slide a nucleosome towards the unoccupied +1 site in R-state. Upon arrival at the +1 site, intrinsic entry-DNA features (A-tracts of NDR) at the +1 site destabilize motor engagement at SHL 6 leading to an off-state conformation. INO80 could still slide the +1 nucleosome back toward the gene body, however, increasing occupancy of the +2 and +3 nucleosomes progressively limits the reverse sliding. Here, shortening linker DNA causes A-module disengagement (A-module flip) and prevalence of an I-state, further restricting movement. When +2 is <30 bp away from +1, further sliding of the +1 nucleosome by INO80 into the gene body is robustly prevented.

**Supplementary Fig. 9:**
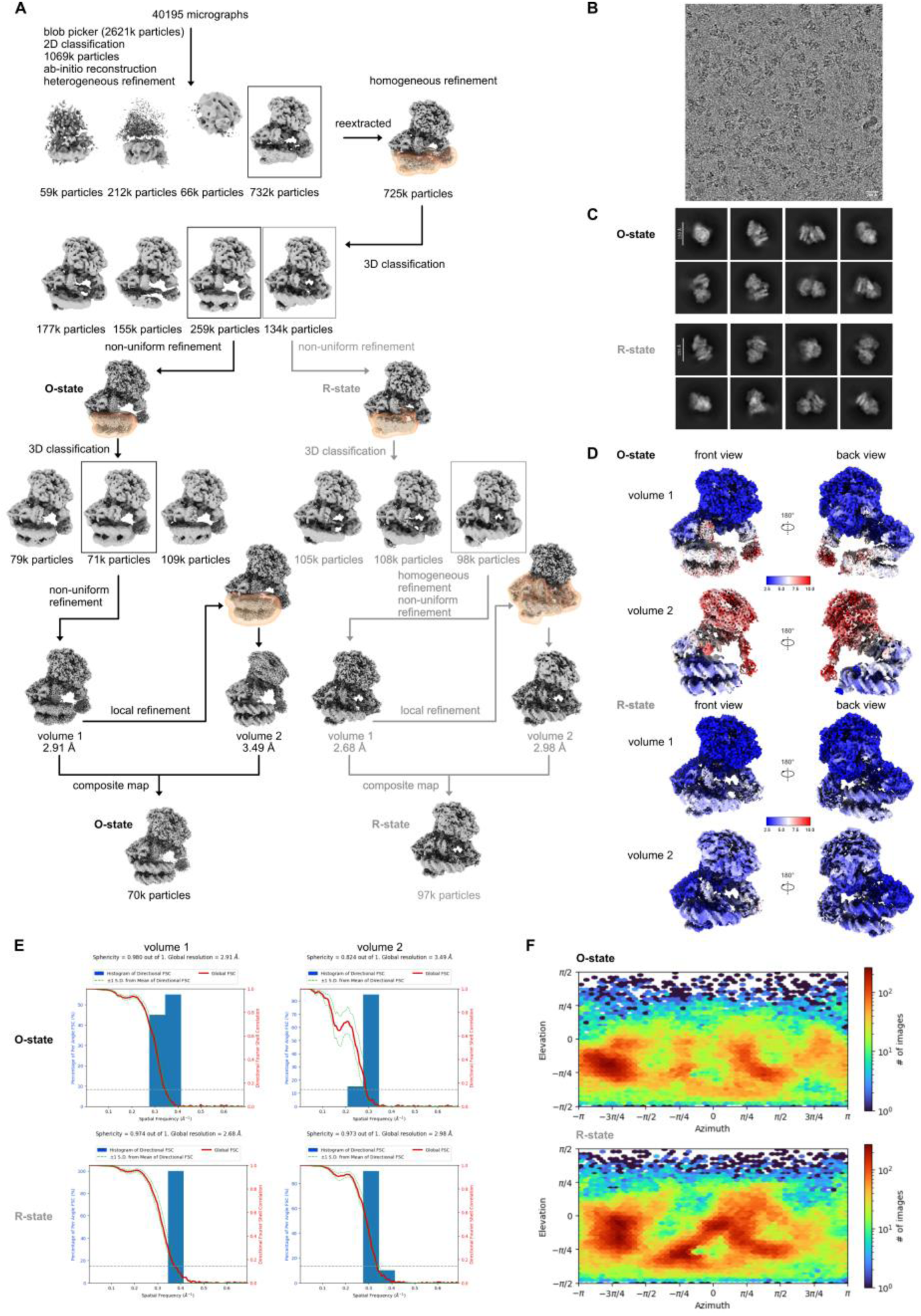
Cryo-EM data analysis of the INO80^ΔN^ C-module bound to array NCP in O- and R-states in the AlF_x_ XL dataset. **A,** Cryo-EM data processing workflow of the INO80^ΔN^-array NCP complexes in O- and R-states performed using cryoSPARC^2^**. B,** Representative micrograph from the 40195 movies collected for the INO80^ΔN^-array NCP in the AlF_x_ XL dataset. **C,** Selected 2D class averages obtained from the particles used in the final reconstruction of the INO80^ΔN^-array NCP complexes in O- and R-states. **D,** Local resolution maps of the INO80^ΔN^-array NCP complex (volume 1 and volume 2) for O- and R-states calculated in cryoSPARC^2^. Regions of higher resolution are shown in blue, lower resolution regions are shown in red. **E,** Fourier shell correlation (FSC) analysis^3^ of volume 1 and volume 2 of the INO80-array NCP complexes in O- and R-states, showing the histogram of directional FSC^2^ (blue) and the global FSC (red) curve. The distribution of directional resolution values is defined as ±1σ (dashed green lines). The grey dashed line marks the 0.143 cutoff criterion, indicating nominal resolutions of 2.91 Å for volume 1 and 3.49 Å for volume 2 in case of O-state and 2.68 Å for volume 1 and 2.98 Å for volume 2 in case of R-state. **F,** Angular distribution of the particles used for the final reconstruction of the INO80-array NCP complexes in O- and R-states in the AlF_x_ XL dataset.

**Supplementary Fig. 10:**
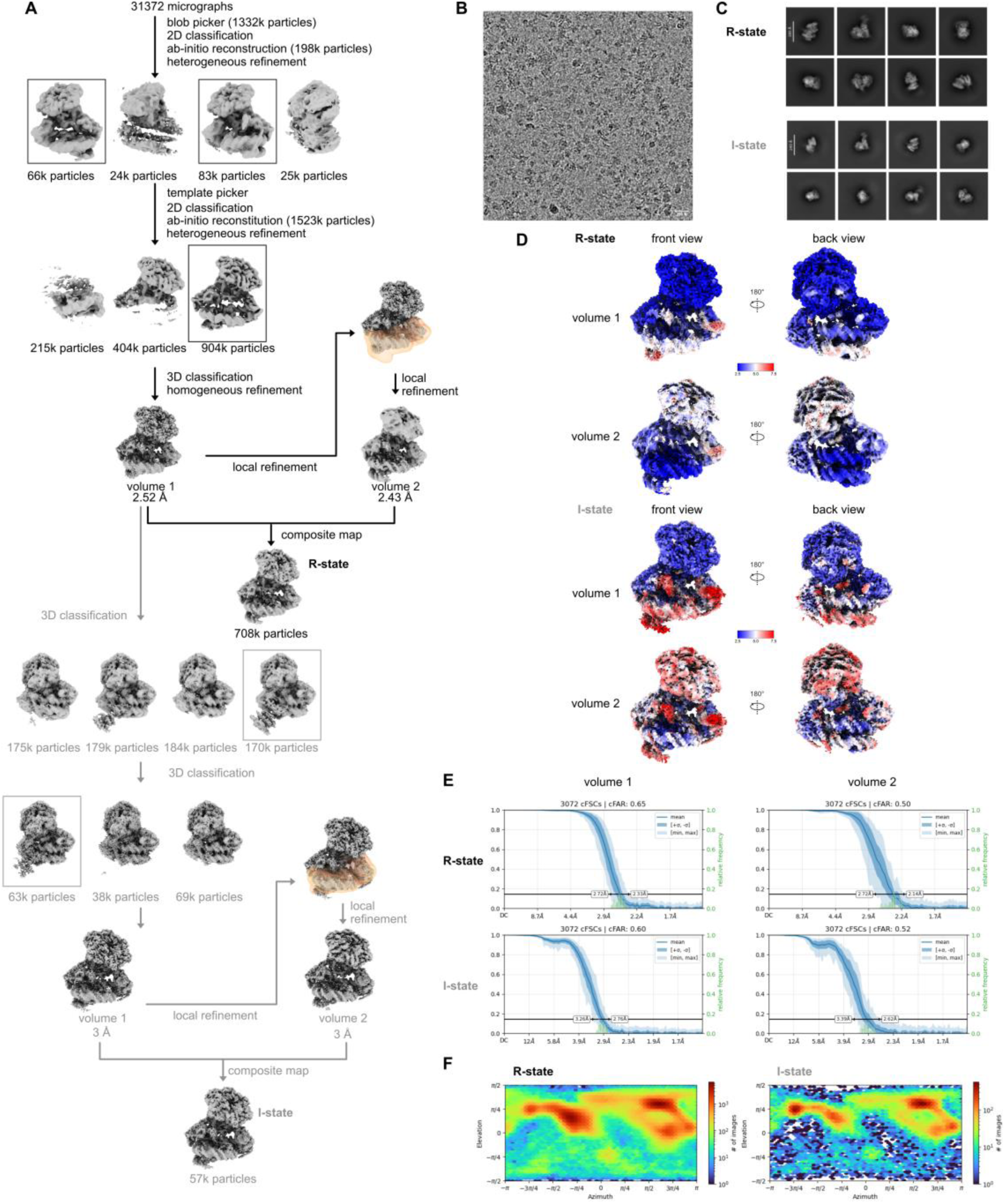
Cryo-EM data analysis of the INO80^ΔN^ C-module bound to array NCP in R- and I-states in the BeF_x_ dataset. **A,** Cryo-EM data processing workflow of the INO80^ΔN^-array NCP complexes in R- and I-states performed using cryoSPARC^2^**. B,** Representative micrograph from the 31372 movies collected for the INO80^ΔN^-array NCP in the BeF_x_ dataset. **C,** Selected 2D class averages obtained from the particles used in the final reconstruction of the INO80^ΔN^-array NCP complexes in R- and I-states. **D,** Local resolution maps of the INO80^ΔN^-array NCP complex (volume 1 and volume 2) for R- and I-states calculated in cryoSPARC^2^. Regions of higher resolution are shown in blue, lower resolution regions are shown in red. **E,** Fourier shell correlation (FSC) analysis^3^ of volume 1 and volume 2 of the INO80-array NCP complexes in R- and I-states, showing the histogram of directional FSC^2^ (blue) and the global FSC (red) curve. The distribution of directional resolution values is defined as ±1σ (dashed green lines). The grey dashed line marks the 0.143 cutoff criterion, indicating nominal resolutions of 2.52 Å for volume 1 and 2.43 Å for volume 2 in case of R-state and 3 Å for volume 1 and 3 Å for volume 2 in case of I-state. **F,** Angular distribution of the particles used for the final reconstruction of the INO80-array NCP complexes in R- and I-states in the BeF_x_ dataset.

**Supplementary Fig. 11:**
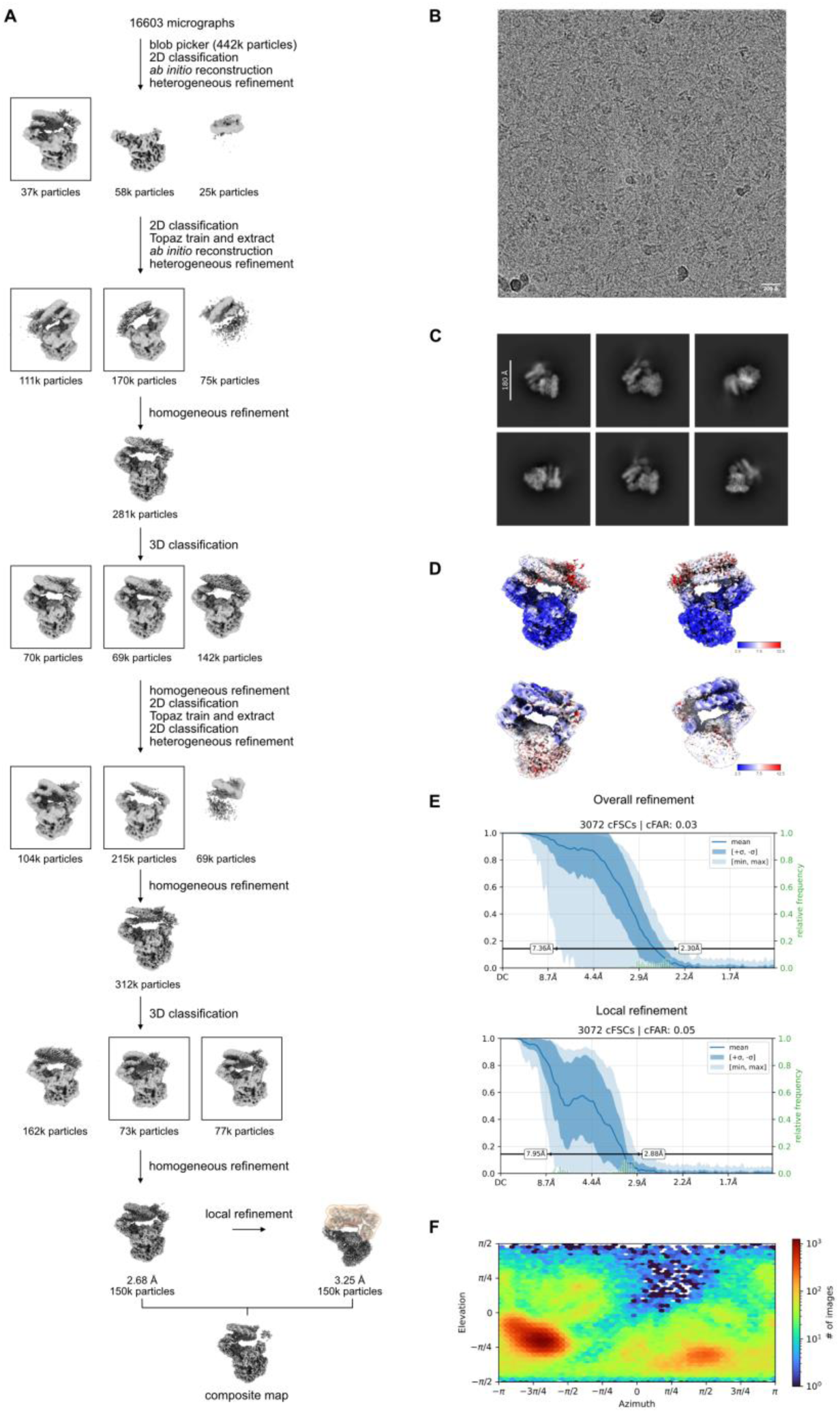
Cryo-EM data analysis of the INO80^ΔN^ C-module bound to the 80S0 nucleosome in the O-state. **A,** Cryo-EM data processing workflow of the INO80^ΔN^-80S0 nucleosome complex performed using cryoSPARC^2^**. B,** Representative micrograph from the 16603 movies collected for the INO80^ΔN^-80S0 nucleosome. **C,** Selected 2D class averages obtained from the particles used in the final reconstruction of the INO80^ΔN^-80S0 nucleosome complex in the O-state. **D,** Local resolution maps of the INO80^ΔN^-80S0 nucleosome complex calculated in cryoSPARC^2^. Regions of higher resolution are shown in blue, lower resolution regions are shown in red. **E,** Conical Fourier shell correlation (FSC) analysis^3^ of homogeneous and local refinements of the INO80^ΔN^-80S0 nucleosome complex states (in blue) and histogram of the spread of resolution values over direction (in green). The black line marks the 0.143 cutoff criterion, indicating nominal resolutions of 2.68 Å for the homogeneous refinement and 3.25 Å for the local refinement. **F,** Angular distribution of the particles used for the final reconstruction of the INO80^ΔN^-80S0 nucleosome complex.

**Supplementary Fig. 12:**
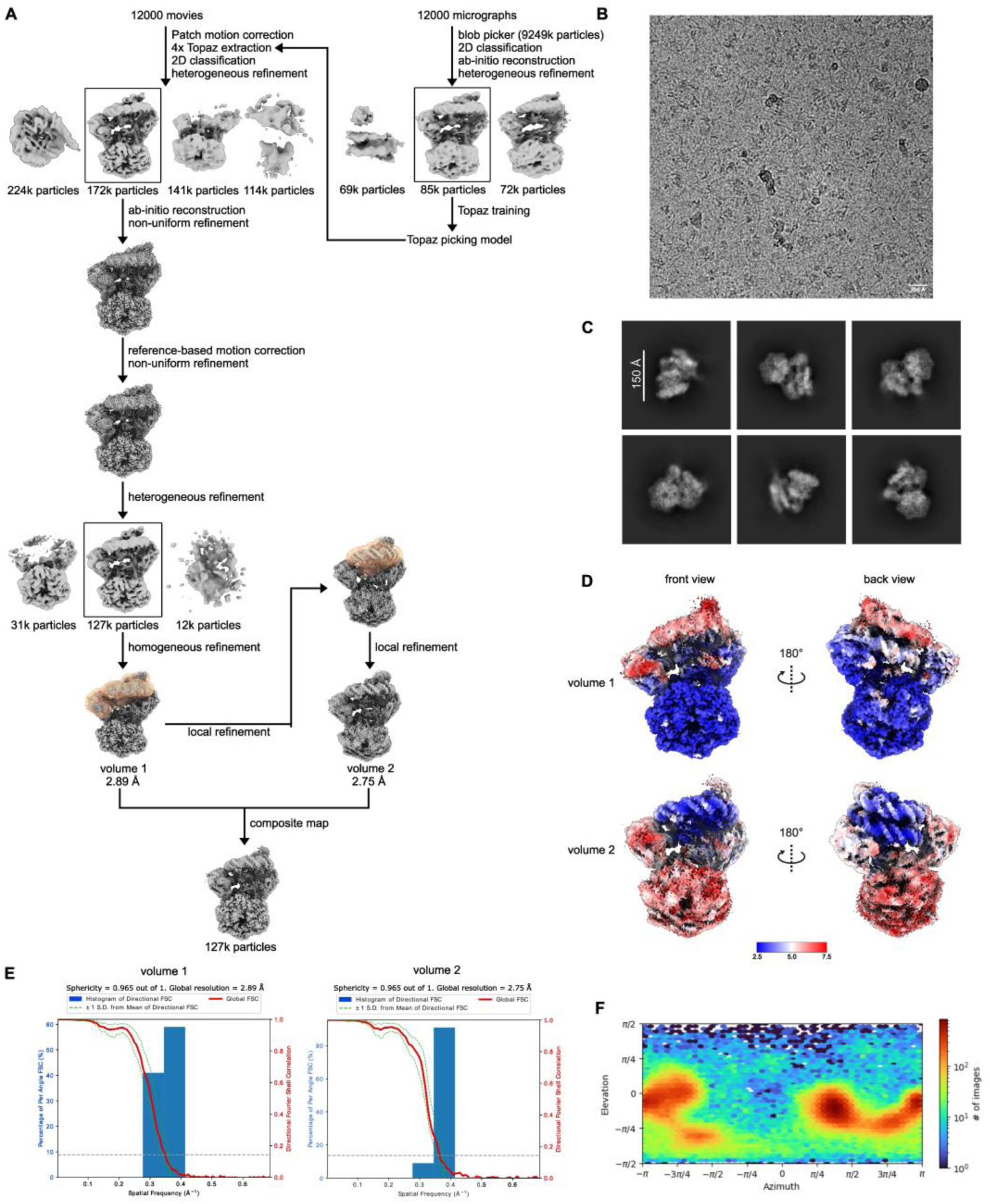
Cryo-EM data analysis of the INO80^fl^-80S80 nucleosome complex. **A,** Cryo-EM data processing workflow of the INO80^fl^-80S80 nucleosome complex performed using cryoSPARC^2^**. B,** Representative micrograph from the 12000 movies collected for the INO80^fl^-80S80 nucleosome dataset. **C,** Selected 2D class averages obtained from the particles used in the final reconstruction of the INO80^fl^-80S80 nucleosome complex. **D,** Local resolution maps of the INO80^fl^- 80S80 nucleosome complex (volume 1 and volume 2) calculated in cryoSPARC^2^. Regions of higher resolution are shown in blue, lower resolution regions are shown in red. **E,** Fourier shell correlation (FSC) analysis^3^ of volume 1 and volume 2 of the INO80^fl^-80S80 nucleosome complex, showing the histogram of directional FSC^2^ (blue) and the global FSC curve (red). The distribution of directional resolution values is defined as ±1σ (dashed green lines). The grey dashed line marks the 0.143 cut-off criterion, indicating nominal resolutions of 2.89 Å for volume 1 and 2.75 Å for volume 2. **F,** Angular distribution of the particles used for the final reconstruction of the INO80^fl^-80S80 nucleosome complex.

**Supplementary Fig. 13:**
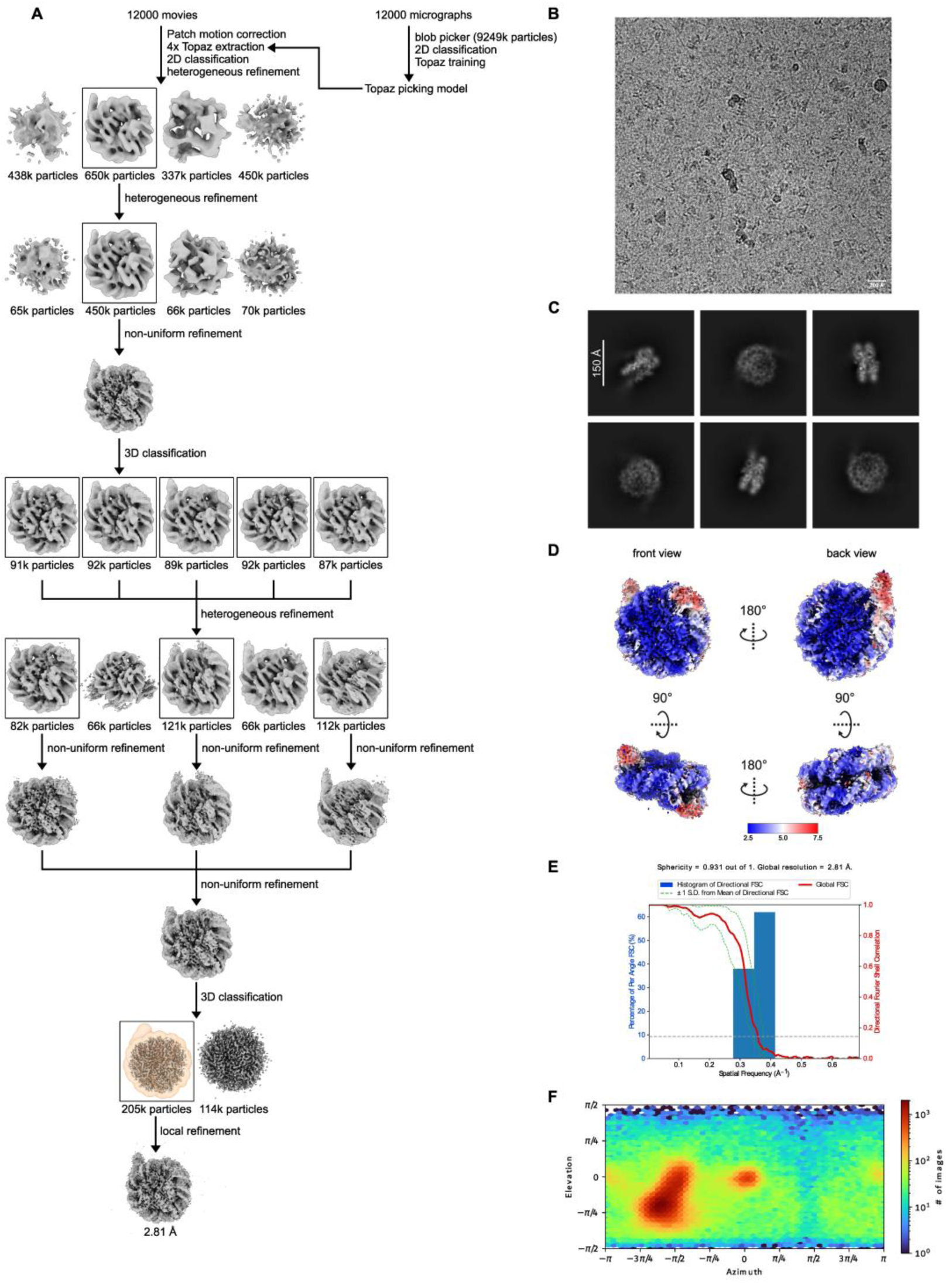
Cryo-EM data analysis of the 80S80 nucleosome. **A,** Cryo-EM data processing workflow of the 80S80 nucleosome performed using cryoSPARC^2^. **B,** Representative micrograph from the 12000 movies collected for the INO80^fl^-80S80 nucleosome dataset. **C,** Selected 2D class averages obtained from the particles used in the final reconstruction of the 80S80 nucleosome. **D,** Local resolution map of the 80S80 nucleosome calculated in cryoSPARC^2^. Regions of higher resolution are shown in blue, lower resolution regions are shown in red. **E,** Fourier shell correlation (FSC) analysis^3^ of the 80S80 final volume, showing the histogram of directional FSC^2^ (blue) and the global FSC (red) curve. The distribution of directional resolution values is defined as ±1σ (dashed green lines). The grey dashed line marks the 0.143 cutoff criterion, indicating nominal resolutions of 2.81 Å. **F,** Angular distribution of the particles used for the final reconstruction of the 80S80 nucleosome.

**Supplementary Fig. 14:**
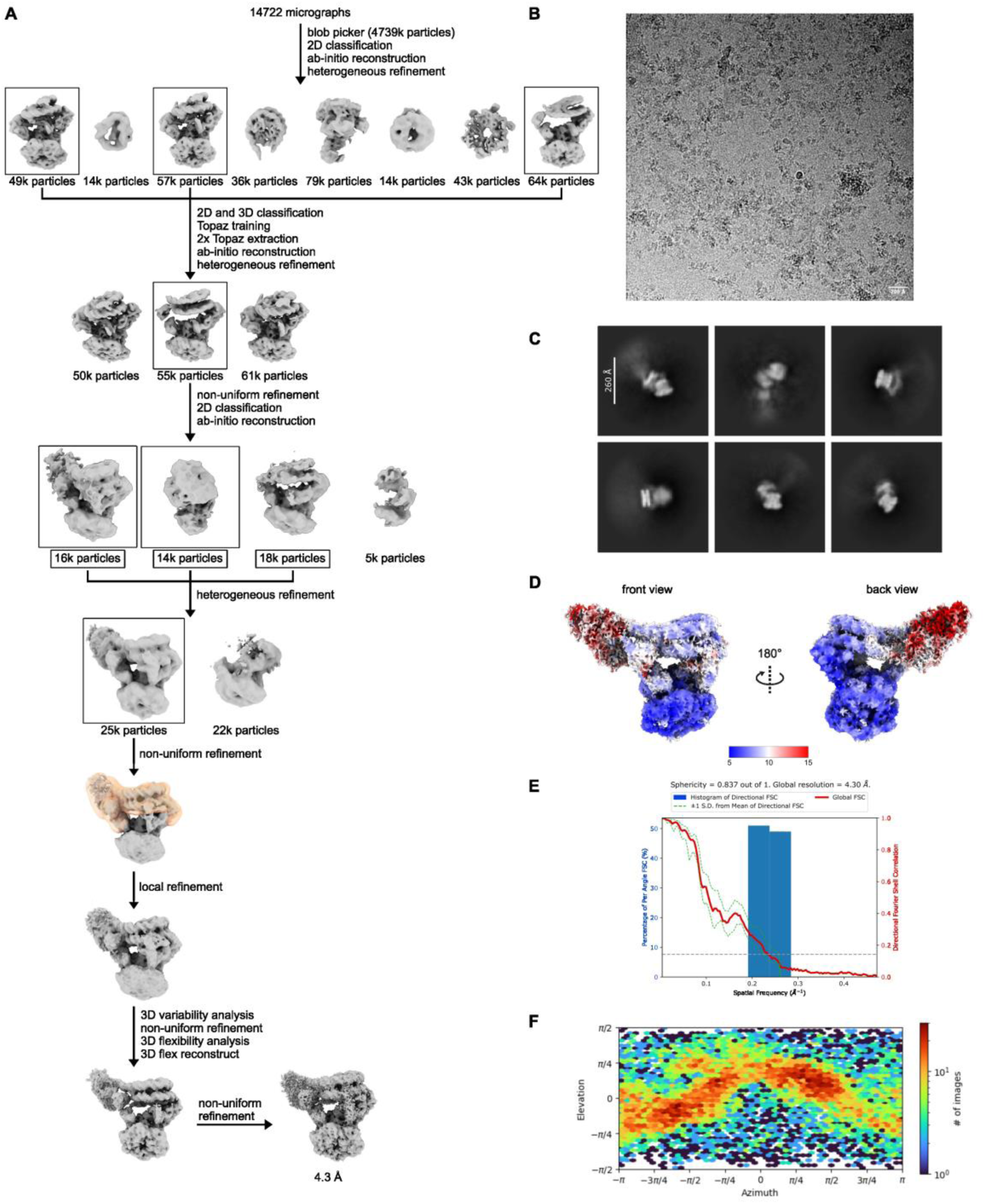
Cryo-EM data analysis of the INO80^fl^-80N0 M2_rev_ nucleosome complex. **A,** Cryo-EM data processing workflow of the INO80^fl^-80N0 M2_rev_ nucleosome complex using cryoSPARC^2^. **B,** Representative micrograph from the 14722 movies collected for the INO80^fl^-80N0 M2_rev_ nucleosome dataset. **C,** Representative 2D class averages obtained from the particles used in the final INO80^fl^-80N0 M2_rev_ nucleosome complex reconstruction. **D,** Visualization of local resolution of the INO80^fl^-80N0 M2_rev_ nucleosome complex calculated in cryoSPARC. Two different orientations are shown (front and back view of the INO80^fl^-80N0 M2_rev_ nucleosome complex). Blue indicates higher resolution, and red indicates lower resolution. **E,** Fourier shell correlation (FSC) analysis^3^ of the INO80^fl^-80N0 M2_rev_ nucleosome complex final volume, showing the histogram of directional FSC (blue) and global FSC curve (red). The spread of directional resolution values is defined as ±1σ (dashed green lines). The grey dashed line indicates the 0.143 cutoff criterion, indicating a nominal resolution of 4.37 Å. **F,** Angular distribution of the particles used for the final 80N0 M2_rev_ nucleosome reconstruction.

**Video S1. Morph of the INO80-nucleosome complex from R-to O-state.**

The animation depicts how INO80 binds to a nucleosome in the R- and O-states. The models are aligned to the histones. ChimeraX visualization of INO80-nucleosome complex morphing from R-state to the sliding-incapable O-state highlights INO80 core flip without full dissociation from the nucleosome. The movie only serves for purposes of illustration and might not reflect the real trajectory of INO80’s movement.

**Table S1.**
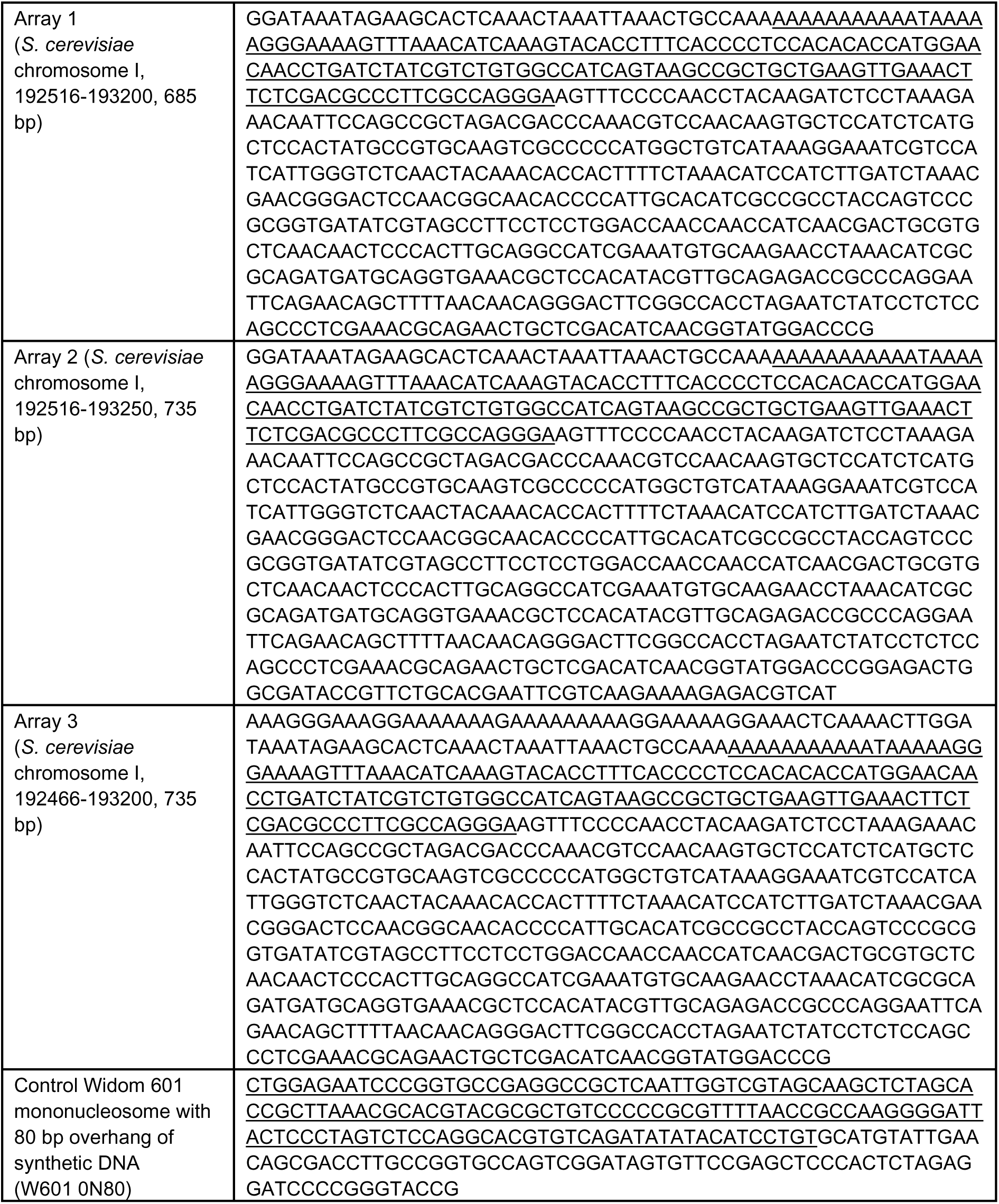
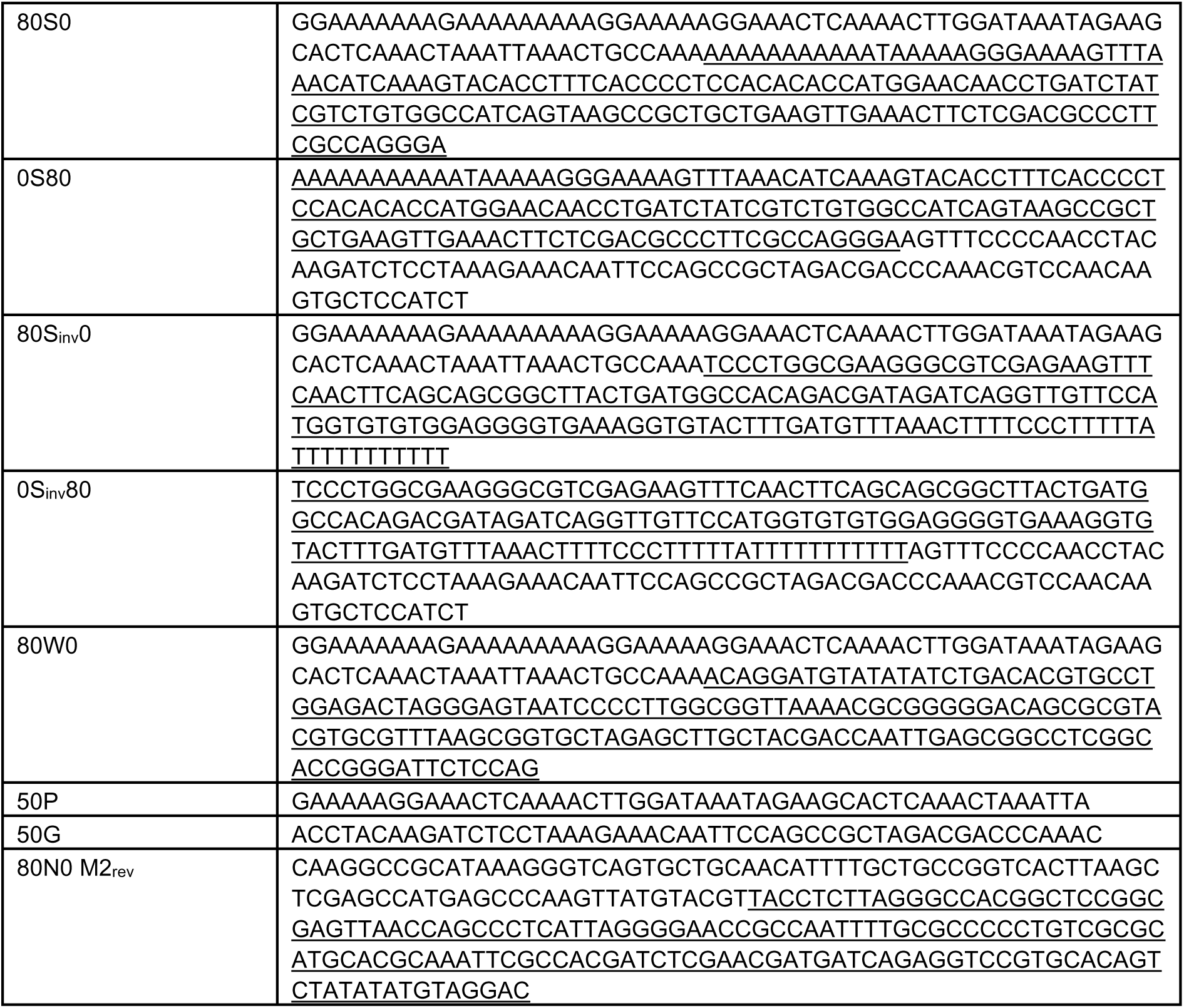
Sequences used in the study (intranucleosomal sequence is underlined where applicable)

**Table S2.** Cryo-EM data collection, refinement and validation statistics for INO80 core bound to array NCP in O-state and INO80 core bound to 80S80 *SWH1* +1 NCP in R-state.

|  | INO80 core bound to array NCP in O-state<br>(EMD-xxx)<br>(PDB xxxx) | INO80 core bound to 80S80 SWH1 +1 NCP in R-state<br>(EMD-xxx)<br>(PDB xxxx) |
| --- | --- | --- |
| **Data collection and processing** |  |  |
| Magnification | 165k | 165k |
| Voltage (kV) | 300 | 300 |
| Electron exposure (e-/Å<sup>2</sup>) | 40 | 40 |
| Defocus range (µm) | -0.5 to -2.6 | -0.5 to -2.6 |
| Pixel size (Å) | 0.727 | 0.727 |
| Symmetry imposed | C1 | C1 |
| Initial particle images (no.) | 2,621,000 | 9,249,000 |
| Final particle images (no.) | 69,872 | 127,169 |
| Map resolution (Å) | 2.91 | 2.75 |
| FSC threshold | 0.143 | 0.143 |
| Map resolution range (Å) | 2.5 to 7.2 | 1.6 to 6.9 |
| **Refinement** |  |  |
| Initial model used (PDB code) | AlphaFold2, 6FML | AlphaFold2, 6WZ5 |
| Model resolution (Å) | 3.0 | 3.0 |
| FSC threshold | 0.143 | 0.143 |
| Map sharpening *B* factor (Å<sup>2</sup>) | 56.6 (see EMD-xxx)<br>52.9 (see EMD-xxx) | 62.9 (see EMD-xxx)<br>68.4 (see EMD-xxx) |
| **Model composition** |  |  |
| Non-hydrogen atoms | 40499 | 42622 |
| Protein residues | 4403 | 4656 |
| Ligands | 7 Mg<sup>2+</sup>, 6 ADP, 1 ATP | 7 Mg<sup>2+</sup>, 3 ADP, 5 ATP |
| Nucleotides | 286 | 282 |
| ***B* factors (Å<sup>2</sup>)** |  |  |
| Protein | 82.3 | 78.1 |
| Ligand | 42.1 | 63.8 |
| **R.m.s. deviations** |  |  |
| Bond lengths (Å) | 0.0138 | 0.0127 |
| Bond angles (°) | 1.83 | 1.88 |
| **Validation** |  |  |
| MolProbity score | 1.67 | 2.01 |
| Clashscore | 4.64 | 6.28 |
| Poor rotamers (%) | 1.35 | 2.34 |
| **Ramachandran plot** |  |  |
| Favored (%) | 95.2 | 94.3 |
| Allowed (%) | 4.7 | 5.2 |
| Disallowed (%) | 0.1 | 0.4 |

**Table S3.** Cryo-EM data collection for INO80 core bound to array NCP in I-state and in R-state (BeF_x_)

| EMDB | Cryo-EM structure of INO80 core bound to array NCP in I-state (BeF<sub>x</sub>) |  |  | Cryo-EM structure of INO80 core bound to array NCP in R-state (BeF<sub>x</sub>) |  |  |
| --- | --- | --- | --- | --- | --- | --- |
|  | overall refinement | nucleosome refinement | composite | overall refinement | nucleosome refinement | composite |
|  | EMD-xxx | EMD-xxx | EMD-xxx | EMD-xxx | EMD-xxx | EMD-xxx |
| **Data collection and processing** |  |  |  |  |  |  |
| Magnification |  |  | 165000x |  |  |  |
| Voltage (kV) |  |  | 300 |  |  |  |
| Electron exposure (e<sup>-</sup>/Å<sup>2</sup>) |  |  | 40 |  |  |  |
| Defocus range (μm) |  |  | -2600 to -1100 |  |  |  |
| Pixel size (Å) |  |  | 0.727 |  |  |  |
| Symmetry imposed |  |  | C1 |  |  |  |
| Final particle images (no.) |  | 56504 |  |  | 707610 |  |
| Map resolution (Å) | 3.00 | 3.00 | 3.00 | 2.52 | 2.43 | 2.43 |
| FSC threshold |  |  | 0.143 |  |  |  |

**Table S4.** Cryo-EM data collection for INO80 core bound to array NCP in R-state (AlF_x_, XL) and for INO80 core bound to *SWH1* +1 NCP in O-state (AlF_x_)

| EMDB | Cryo-EM structure of INO80 core bound to array NCP in R-state (AlF<sub>x</sub>, XL) |  |  | Cryo-EM structure of INO80 core bound to SWH1 +1 NCP in O-state (AlF<sub>x</sub>) |  |  |
| --- | --- | --- | --- | --- | --- | --- |
|  | overall refinement | nucleosome refinement | composite | overall refinement | nucleosome refinement | composite |
|  | EMD-xxx | EMD-xxx | EMD-xxx | EMD-xxx | EMD-xxx | EMD-xxx |
| **Data collection and processing** |  |  |  |  |  |  |
| Magnification |  |  | 165000x |  |  |  |
| Voltage (kV) |  |  | 300 |  |  |  |
| Electron exposure (e<sup>-</sup>/Å<sup>2</sup>) |  |  | 40 |  |  |  |
| Defocus range (μm) |  |  | -2600 to -500 |  |  |  |
| Pixel size (Å) |  |  | 0.727 |  |  |  |
| Symmetry imposed |  |  | C1 |  |  |  |
| Final particle images (no.) |  | 97355 |  |  | 149945 |  |
| Map resolution (Å) | 2.68 | 2.98 | 2.68 | 2.68 | 3.25 | 2.68 |
| FSC threshold |  |  | 0.143 |  |  |  |

**Table S5.**
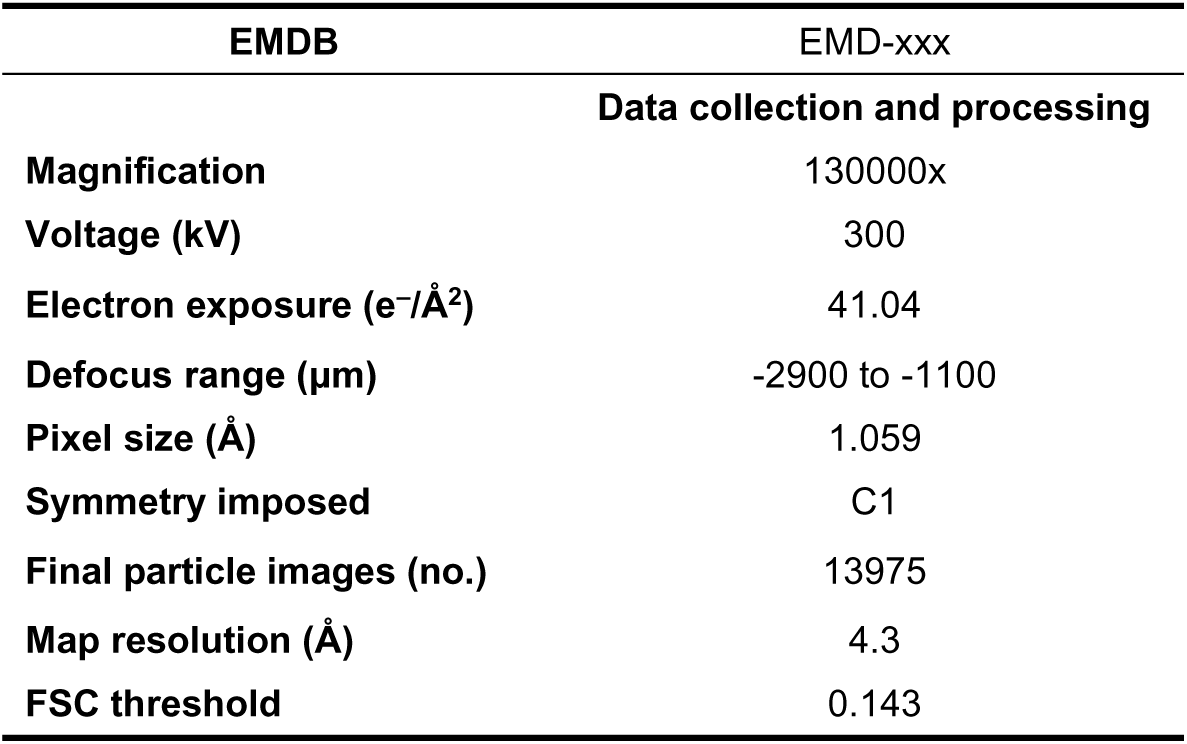
Cryo-EM data collection, refinement and validation statistics for INO80 core and A-module bound to M2_rev_ nucleosome in O-state.

| EMDB | EMD-xxx |
| --- | --- |
| **Data collection and processing** |  |
| Magnification | 130000x |
| Voltage (kV) | 300 |
| Electron exposure (e<sup>-</sup>/Å<sup>2</sup>) | 41.04 |
| Defocus range (µm) | -2900 to -1100 |
| Pixel size (Å) | 1.059 |
| Symmetry imposed | C1 |
| Final particle images (no.) | 13975 |
| Map resolution (Å) | 4.3 |
| FSC threshold | 0.143 |

## Supplementary Method: Scoring DNA Sequences in cryo-EM Maps

### A) Workflow

This procedure aims to identify the most likely nucleic acid (NA) sequence represented in a cryo-EM map. It does so by calculating map-model correlation coefficients for possible bases and base pairs and then identifying the highest-scoring sequence segment within a target sequence using a sliding-window approach. Determining the identity of individual bases or base pairs is typically very difficult because of map-resolution limitations. However, collective analysis of multiple consecutive bases or base pairs in an NA chain increases statistical power, provided that the candidate target NA sequence is known and the sequence space is not too large. Our pipeline follows the general procedure outlined previously^1^. Rather than using a neural-network-based approach to identify bases or purine/pyrimidine classes, it uses map-model correlation coefficients derived from established structural biology software.

The computational workflow is as follows:

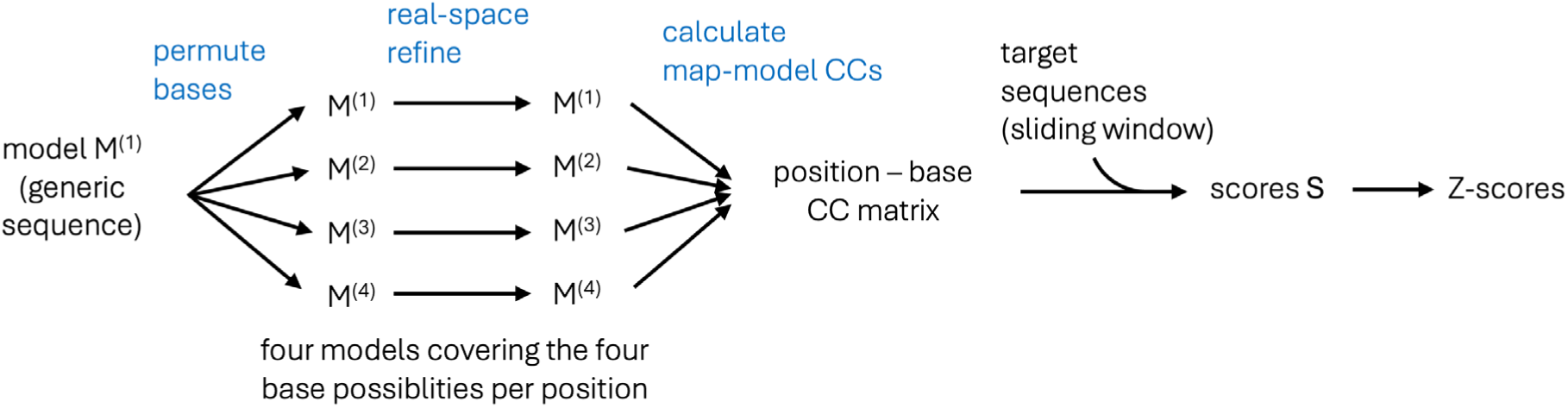

(i) Starting from a user-provided atomic model M^(1)^ containing nucleic acid N^(1)^, three additional models, M^(2)^-M^(4)^, with nucleic acids N^(2)^-N^(4)^ are generated. The non-nucleic acid parts of the model remain unchanged, whereas the bases in the nucleic acid are altered so that all four possible base types (A, T, G, and C) are sampled at each base position. To generate all four possible base types at each position while preserving any Watson–Crick base pairing present in the starting model, the nucleotides are permuted using ChimeraX^2^ `swapna` in the following way:
(ii)

N^(2)^ = π_2_(N^(1)^), with π_2_: A→G, T→C, G→A, C→T

N^(3)^ = π_3_(N^(1)^), with π_3_: A→T, T→A, G→C, C→G

N^(4)^ = π_4_(N^(1)^), with π_4_: A→C, T→G, G→T, C→A

(iii) Each model M^(m)^, where m ∈ {1,2,3,4}, is then real-space refined against the map. In our analysis, we use 50 cycles of Servalcat^3^ with jelly-body constraints.
(iv) Correlation coefficients, cc_im_, are then calculated to form a position-base matrix, where i ∈ {1, …, n} denotes the base position and m ∈ {1, 2, 3, 4} denotes the model. The cc_im_ values are obtained using Phenix^4^ `phenix.map_correlations` on models M^(m)^.
(v) Optionally, to include information on base pairing, we can use the root mean square average of the correlation coefficients for base-paired positions i and k:

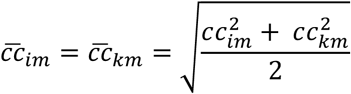

(vi) With the cc_im_ values, or the corresponding paired values cc_im_, we can construct a position-base matrix CC ∈ ℝ*^i^*^4^ with elements cc_ib_, where i ∈ {1,…,n} and b ∈ {A,T,G,C}). From this matrix, we calculate an overall score S(T) for a target sequence T:

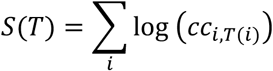

where T(i) denotes the base type at position i in T. Alternative scoring functions include those that also account for the complement T* of T:

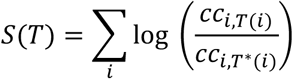

or position-wise normalized cc_im_ values:

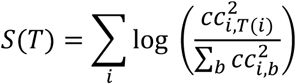

Additional weighting factors may include Q-scores (resolvability) or cutoff filters to exclude poorly fitting positions in the map, for example positions with low correlation coefficients. Overall, the different scoring functions gave qualitatively similar results, with only small quantitative differences. Which function yielded higher Z-scores depended on the dataset, and a broader benchmark analysis will be required to determine which performs best across a large set of cases.

(vii) Typically, the goal is to determine whether the map is consistent with a particular segment of a longer target NA sequence used in the experiment. To analyze the mean, variance, and highest-scoring positions in a target sequence that is longer than the sequence defined by the map/model, we use a sliding-segment approach and calculate a Z-score for segment T_j_ within T, where the length of T_j_ corresponds to the length of the sequence defined by the model:

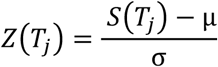

Here, S(T_j_) is the score calculated for a particular segment, and μ and σ denote the mean and standard deviation of the score distribution, respectively. To increase the number of background sequences used for Z-score calculation, the target sequence T can be padded with random permutations of T so as to preserve base composition.

### B) Tests and Examples

In this section, we provide test cases to validate the approach.

#### 1) hOGG1-nucleosome complex (EMD-19870)

Nucleosomes exhibit approximate pseudo-twofold symmetry: they are symmetric with respect to the protein core and DNA backbone, but asymmetric with respect to base identity. These subtle asymmetric features pose a challenge for cryo-EM processing algorithms attempting to determine particle orientation unambiguously. As a result, the reconstructed map can represent a superposition of two orientations related by pseudo-twofold symmetry. To test our method, we required an example containing additional asymmetric features that would allow cryo-EM processing to determine the nucleosome orientation unambiguously. We therefore used a 3.1 Å structure of a nucleosome containing a single 8-oxoguanine (8-oxoG) lesion at SHL 6. In this structure, 8-oxoG is specifically recognized by human 8-oxoguanine DNA glycosylase 1 (hOGG1), giving rise to a well-defined hOGG1–nucleosome complex^5^ (**Fig. 1a**).

Applying the above procedure to a padded Widom-601^6^ sequence (strand 1) produced a single peak (Z-score ∼10) for one chain (Y) of the model, but not for the other chain (Z) (**Fig. 1b**). Using the reverse complement (strand 2) as the target produced a corresponding peak for chain Z, but not for chain Y. Minor peaks (Z-scores ∼3) for strand 1 matching chain Z could reflect a minor population of nucleosomes in the opposite orientation, perhaps arising from contamination by nucleosomes not bound by hOGG1. In any case, the locations of these two peaks correspond to the two strands of a base-paired DNA duplex. The resulting sequence assignment and orientation place the 8-oxoG lesion in the active site of hOGG1 and are consistent with the established Widom-601 positioning around the nucleosome.

**Figure 1:**
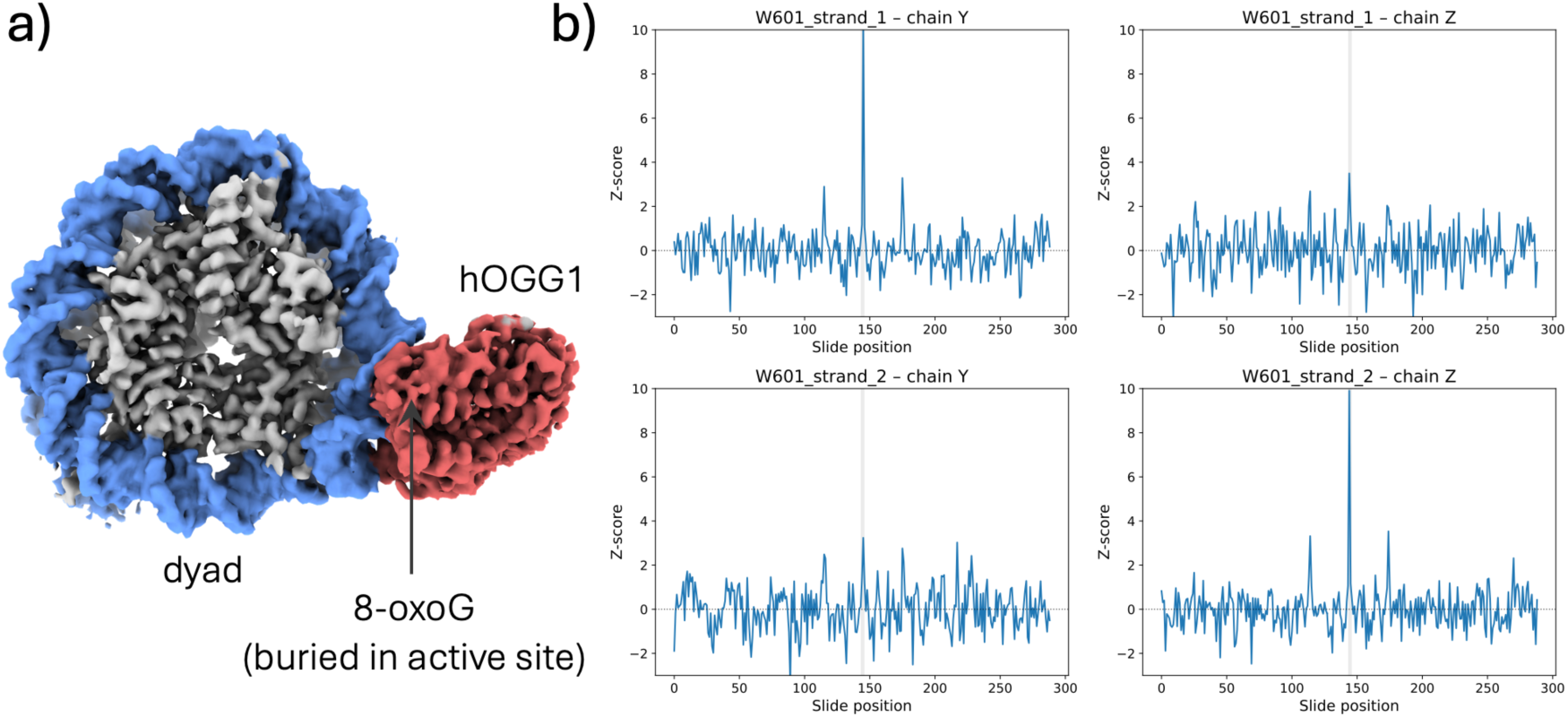
a) 3.1 Å cryo-EM map of DNA glycosylase hOGG1 bound to a nucleosome containing a single 8-oxoG lesion (EMD-19870). b) The nucleic acid mapping procedure identifies a distinct peak for Widom-601 strand 1 in chain Y of the PDB file, and an equivalent distinct peak for the reverse complement of Widom-601 (strand 2) in chain Z. The thin grey region indicates the sliding positions for which the query segment lies entirely within the target sequence and not in the padded flanking regions.

#### 2) hOGG1-nucleosome complex (EMD-43597)

An independent study yielded a similar hOGG1–nucleosome complex^7^, (**Fig. 2a**), again containing a single 8-oxoG lesion at SHL 6, but in the opposite strand relative to EMD-19870. The resulting map-correlation analysis again revealed a single peak (Z-score ∼10) for strand 1 of the padded Widom-601 sequence in one DNA chain of the PDB file, and a single peak for strand 2 in the other chain (**Fig. 2b**). As in the previous example, the resulting sequence assignment places the 8-oxoG lesion in the active site of hOGG1.

**Figure 2:**
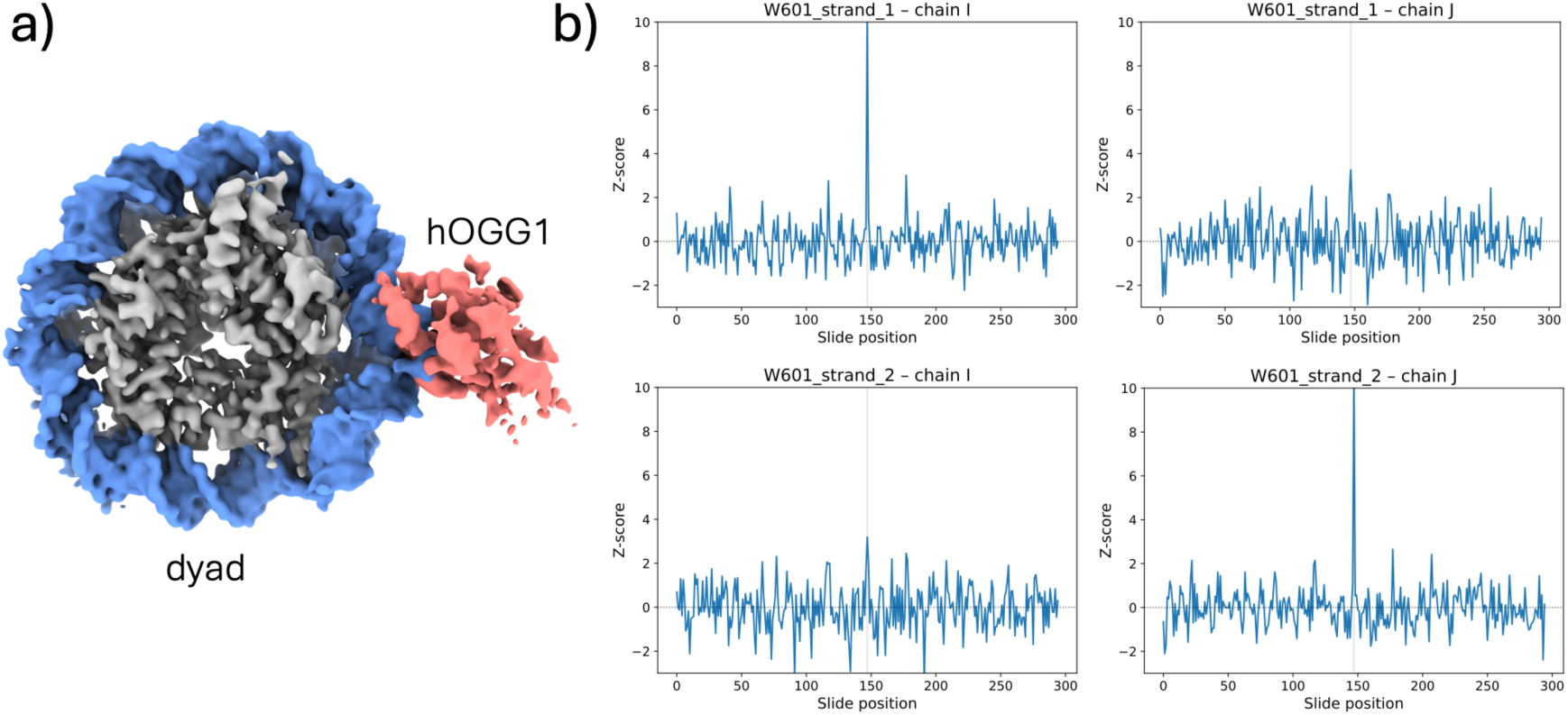
a) 3.3 Å cryo-EM map of DNA glycosylase hOGG1 bound to a nucleosome containing a single 8-oxoG lesion (EMD-43597). b) The nucleic acid mapping procedure identifies a single peak for Widom-601 strand 1 in chain I of the PDB file, and a single peak for the reverse complement of Widom-601 (strand 2) in chain J

#### 3) Unbound 8-oxoG nucleosomes (EMD-43600)

To test whether two superposed sequence assignments can be identified, we also applied our map-correlation pipeline to the map and model of the unbound nucleosome from the same hOGG1 study^7^ (**Fig. 3a**). In this case, distinct peaks were observed for strand 1 in both DNA chains I and J in the coordinate file, and likewise for strand 2 (**Fig. 3b**). This is consistent with the idea that the cryo-EM processing could not resolve the small asymmetric features in these nucleosome particles, resulting in two superposed orientations. Importantly, however, given sufficient map quality, the pipeline can distinguish the two superposed sequence assignments.

**Figure 3:**
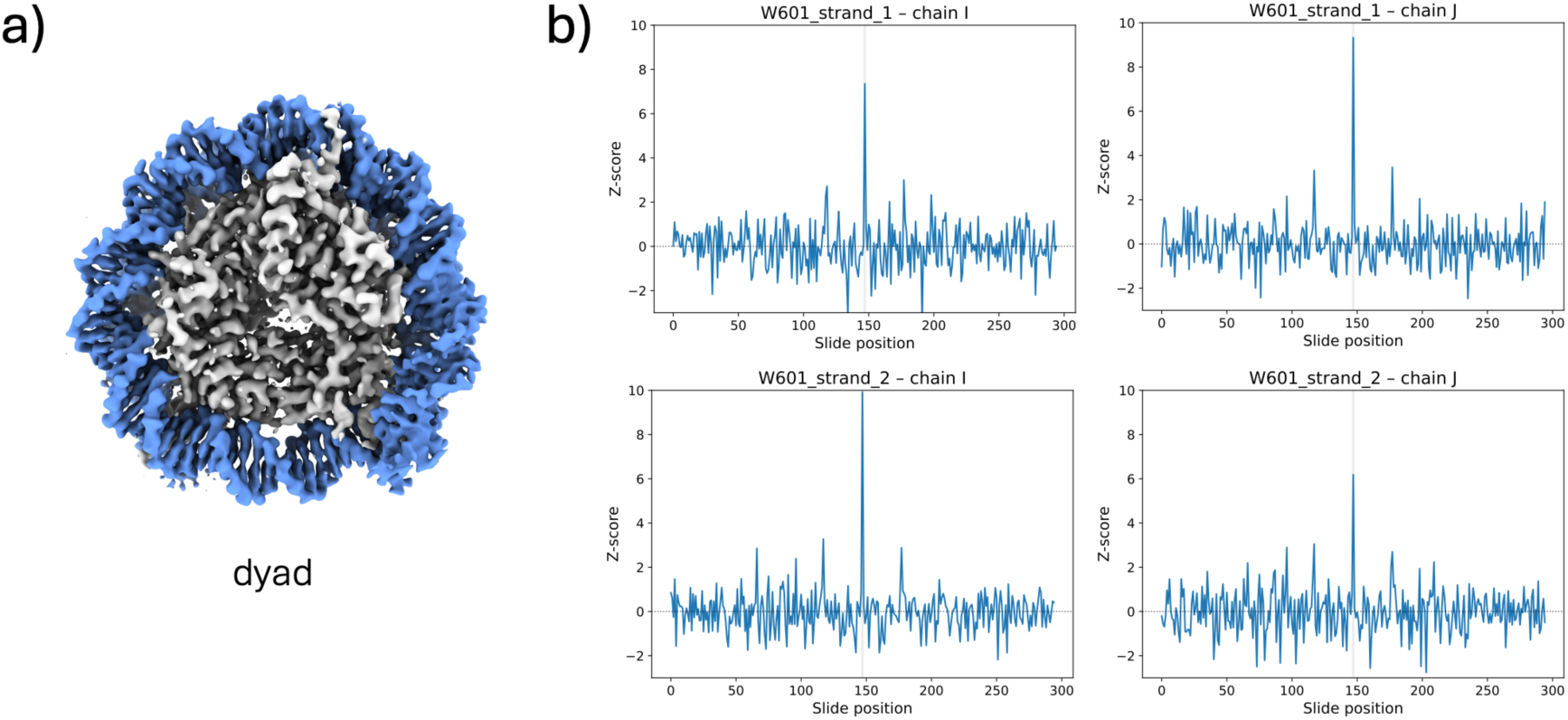
a) 3.0 Å cryo-EM map of the free 8-oxoG nucleosome (EMD-43600). b) The nucleic acid mapping procedure identifies peaks for both strands of the Widom-601 sequence in both chains I and J.

#### 4) hOGG1-nucleosome complex with imposed C2 symmetry

To further test whether the method can deconvolute two superposed sequence assignments, we reprocessed the asymmetric hOGG1–nucleosome complex particles from EMD-19870 with enforced twofold symmetry (**Fig. 4a**). As expected, prominent Z-score matches were now obtained for both strands 1 and 2 in chain Y, and similarly for chain Z (**Fig. 4b**). This analysis demonstrates that the approach can identify at least two superposed and averaged sequence assignments, provided that map quality is sufficient.

**Figure 4:**
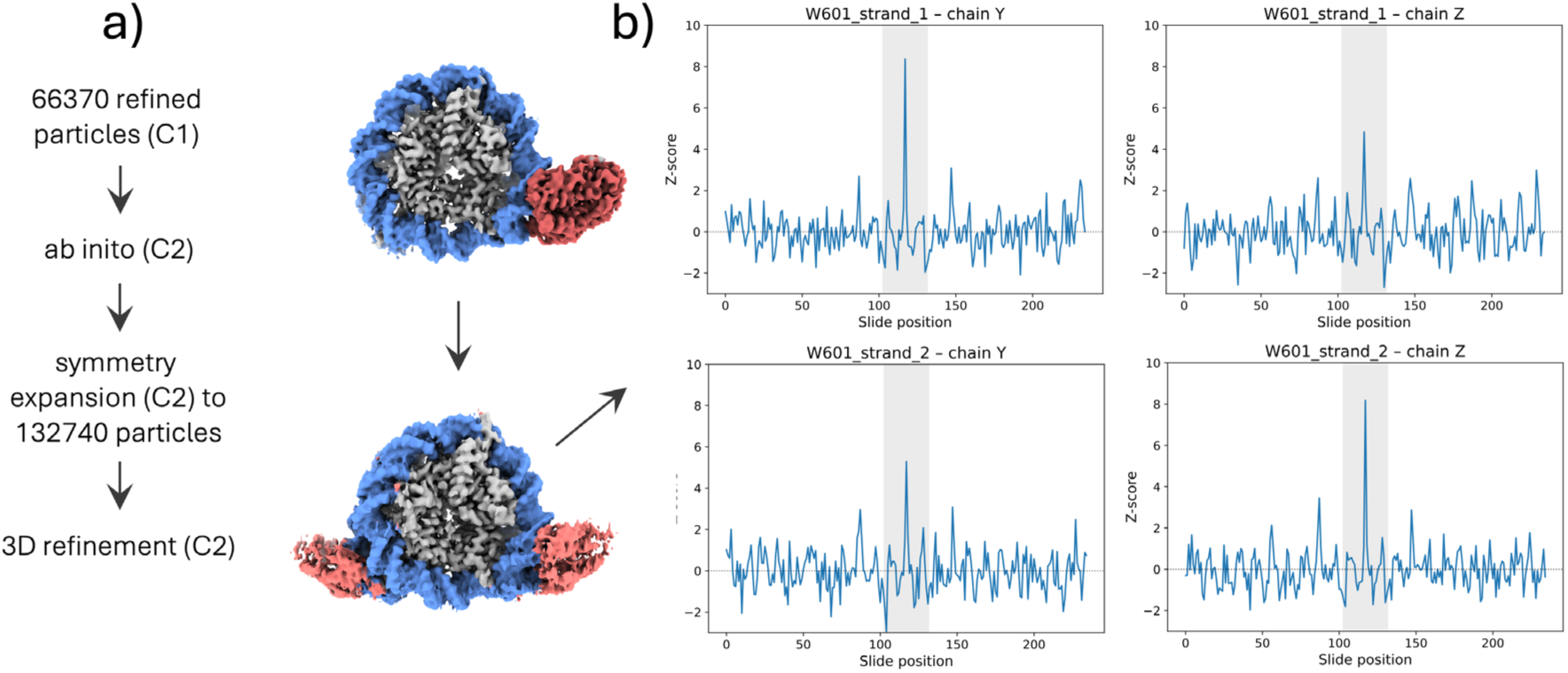
a) 3.0 Å cryo-EM map of the hOGG1–nucleosome complex reprocessed with imposed C2 symmetry. b) The nucleic acid mapping procedure identifies peaks for both strands of the Widom-601 sequence in both chains Y and Z.

#### 5) S. cerevisiae INO80-hexasome complex (EMD-28602)

A hexasome can be assembled on a Widom-601 sequence in a defined orientation; that is, the H2A/H2B dimer is preferentially missing from one side of the asymmetric Widom-601 nucleosome^8^. Such a hexasome substrate, with additional extranucleosomal DNA attached on the H2A/H2B-deficient side, was used to determine a 2.9 Å focused map of the hexasome in complex with INO80^9^ **(Fig. 5a)**. Our pipeline revealed a distinct match corresponding to the proposed DNA assembly on the hexasome in an end-positioned configuration, as well as the expected orientation of the hexasome within the INO80 complex **(Fig. 5b)**.

**Figure 5:**
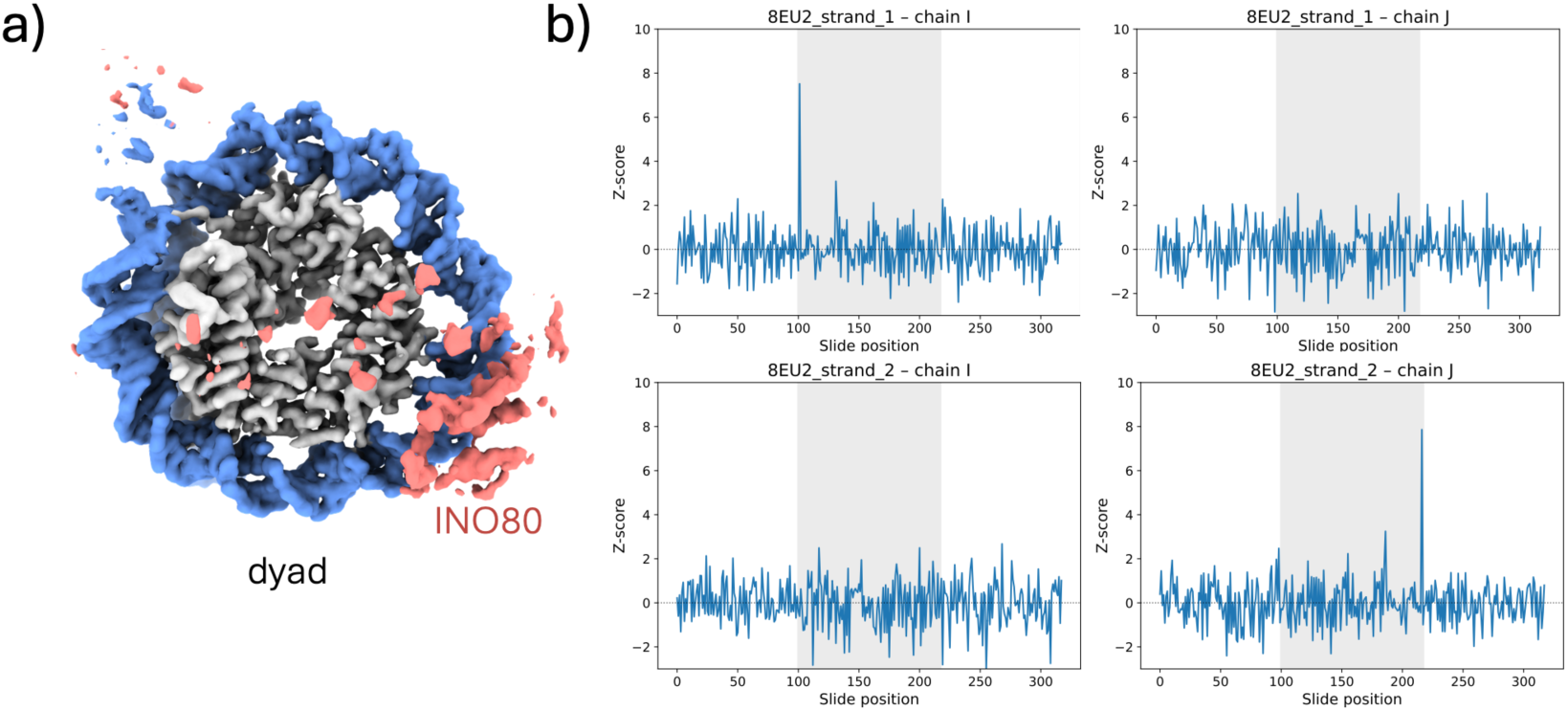
a) 2.9 Å cryo-EM map of the focused-refined hexasome bound to yeast INO80 (EMD-28602). b) The nucleic acid mapping procedure identifies matching peaks for strand 1 in chain I and for strand 2 in chain J of the model.

#### 6) Centromeric nucleosome assembled on native alpha satellite DNA (EMD-0586)

To validate the approach on a non-Widom-601 sequence, we analyzed the 3.4 Å map and model of a centromeric (CENP-A) nucleosome assembled with alpha satellite DNA^10^ (**Fig. 6a**). We again observed distinct high Z-score peaks that match the established sequence register (**Fig. 6b**).

**Figure 6:**
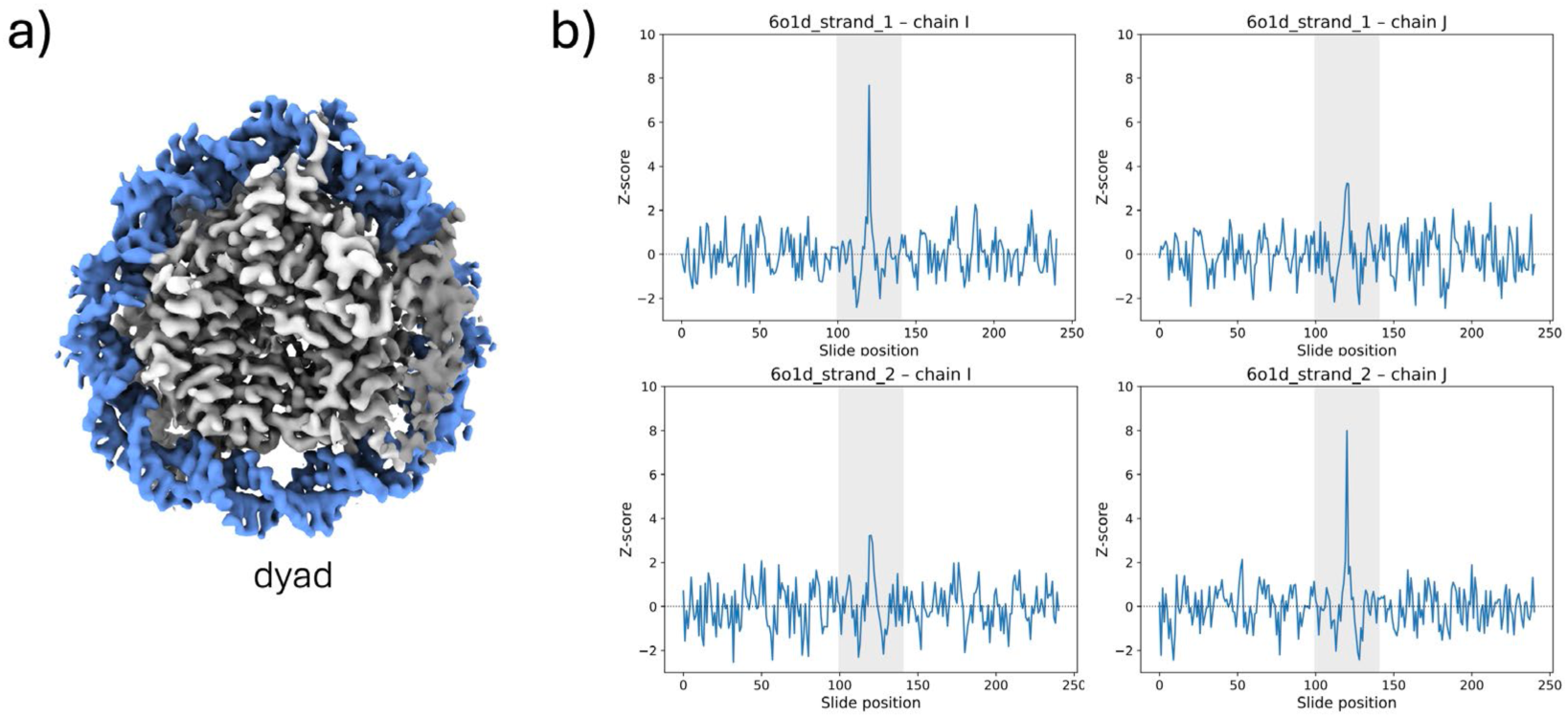
a) 3.4 Å cryo-EM map of the CENP-A nucleosome assembled on alpha satellite DNA (EMD-0586). The grey portion of the DNA density was excluded from the analysis because of poor map quality. b) The nucleic acid mapping procedure correctly identifies matching peaks for strand 1 in chain I and strand 2 in chain J of the model. Smaller peaks are also observed for the orientation related by pseudo-twofold symmetry, suggesting that the map is a superposition of a minor and a dominant orientation.

#### 7) Semi-synthetic data

To further assess the extent to which the approach can resolve multiple superposed sequence assignments, we used semi-synthetic data. We began with the cc values derived from EMD-19870 (see example 1) and identify, at each map position i, the sets (cc_i_^A^, cc_i_^T^, cc_i_^G^, cc_i_^C^)^bi^, where b_i_ ∈ {A,T,G,C} denotes the correct base at position i as inferred from the EMD-19870 analysis. This provides a reasonable estimate of the ground truth, based on the results obtained for that dataset. For each base type b ∈ {A,T,G,C}, we therefore obtain a collection of (cc_i_^A^, cc_i_^T^, cc_i_^G^, cc_i_^C^)^b^ to draw from which synthetic map model correlations for a sequence N^syn^ could be sampled.

To simulate a superposition of k sequences N_k_, we generate for each base position i in N_k_ the set of cc values (cc_ik_^A^, cc_ik_^T^, cc_ik_^G^, cc_ik_^C^)^b_ik^. A simulated map model correlation matrix obtained from “superposed” sequences N_k_ may be in a first approximation constructed as:

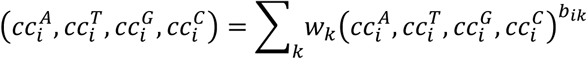

with ∑*_k_w_k_*=1 and an element-wise summation. It was important, however, to sort (cc_i_^A^, cc_i_^T^, cc_i_^G^, cc_i_^C^)^b_i^ into bins based on the average cc values in this set and then combine (cc_i_^A^, cc_i_^T^, cc_i_^G^, cc_i_^C^)^b_ik^ from the same or adjacent bin. This was necessary because the differences between individual cc_i_^b^ values in each positional set can be smaller than differences between corresponding cc_i_^b^ values from different sets i. Using this procedure, we analyzed a range of simulated superposed sequences, as though the nucleosome could assemble at different positions along a padded Widom-601 sequence. Although this simplified model may overestimate resolvability, we were able to identify up to five superposed sequences.

**Figure 7:**
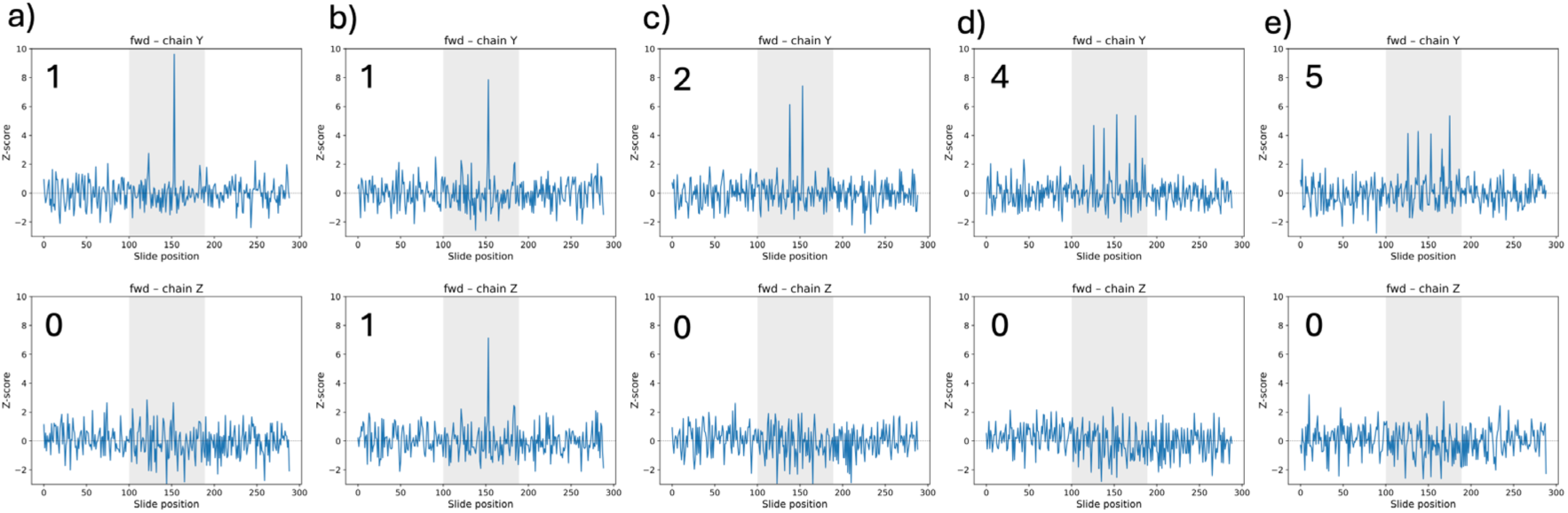
Examples of simulated superposed maps. a), c), d), and e) are simulated averages containing 1, 2, 4, and 5 superposed positions of the forward target strand, respectively. These are correctly identified by distinct high Z-score values for chain Y, but not chain Z, of the model. b) shows a simulated average of superposed forward and reverse-complement strands, which are correctly identified in both chain Y and chain Z of the model.

